# Critical Assessment of Metagenome Interpretation - the second round of challenges

**DOI:** 10.1101/2021.07.12.451567

**Authors:** F. Meyer, A. Fritz, Z.-L. Deng, D. Koslicki, A. Gurevich, G. Robertson, M. Alser, D. Antipov, F. Beghini, D. Bertrand, J. J. Brito, C.T. Brown, J. Buchmann, A. Buluç, B. Chen, R. Chikhi, P. T. Clausen, A. Cristian, P. W. Dabrowski, A. E. Darling, R. Egan, E. Eskin, E. Georganas, E. Goltsman, M. A. Gray, L. H. Hansen, S. Hofmeyr, P. Huang, L. Irber, H. Jia, T. S. Jørgensen, S. D. Kieser, T. Klemetsen, A. Kola, M. Kolmogorov, A. Korobeynikov, J. Kwan, N. LaPierre, C. Lemaitre, C. Li, A. Limasset, F. Malcher-Miranda, S. Mangul, V. R. Marcelino, C. Marchet, P. Marijon, D. Meleshko, D. R. Mende, A. Milanese, N. Nagarajan, J. Nissen, S. Nurk, L. Oliker, L. Paoli, P. Peterlongo, V. C. Piro, J. S. Porter, S. Rasmussen, E. R. Rees, K. Reinert, B. Renard, E. M. Robertsen, G. L. Rosen, H.-J. Ruscheweyh, V. Sarwal, N. Segata, E. Seiler, L. Shi, F. Sun, S. Sunagawa, S. J. Sørensen, A. Thomas, C. Tong, M. Trajkovski, J. Tremblay, G. Uritskiy, R. Vicedomini, Zi. Wang, Zhe. Wang, Zho. Wang, A. Warren, N. P. Willassen, K. Yelick, R. You, G. Zeller, Z. Zhao, S. Zhu, J. Zhu, R. Garrido-Oter, P. Gastmeier, S. Hacquard, S. Häußler, A. Khaledi, F. Maechler, F. Mesny, S. Radutoiu, P. Schulze-Lefert, N. Smit, T. Strowig, A. Bremges, A. Sczyrba, A. C. McHardy

## Abstract

Evaluating metagenomic software is key for optimizing metagenome interpretation and focus of the community-driven initiative for the Critical Assessment of Metagenome Interpretation (CAMI). In its second challenge, CAMI engaged the community to assess their methods on realistic and complex metagenomic datasets with long and short reads, created from ∼1,700 novel and known microbial genomes, as well as ∼600 novel plasmids and viruses. Altogether 5,002 results by 76 program versions were analyzed, representing a 22x increase in results.

Substantial improvements were seen in metagenome assembly, some due to using long-read data. The presence of related strains still was challenging for assembly and genome binning, as was assembly quality for the latter. Taxon profilers demonstrated a marked maturation, with taxon profilers and binners excelling at higher bacterial taxonomic ranks, but underperforming for viruses and archaea. Assessment of clinical pathogen detection techniques revealed a need to improve reproducibility. Analysis of program runtimes and memory usage identified highly efficient programs, including some top performers with other metrics. The CAMI II results identify current challenges, but also guide researchers in selecting methods for specific analyses.

## Introduction

Over the last two decades, advances in metagenomic techniques have vastly increased our knowledge of the microbial world and intensified development of data analysis techniques^1–5^. This created a need for unbiased and comprehensive performance assessment of these methods, to identify best practices as well as open challenges in the field^6–13^. CAMI, the Initiative for the Critical Assessment of Metagenome Interpretation, is a community-driven effort that addresses this need, by offering comprehensive benchmarking challenges and datasets representing common experimental settings, data generation techniques, and environments in microbiome research. In addition to its open and collaborative nature, data FAIRness and reproducibility are key defining principles^14^.

The first CAMI challenge delivered insights into the performances of metagenome assembly, genome and taxonomic binning and profiling programs across multiple complex benchmark datasets, including unpublished genomes across a range of evolutionary divergences and of poorly categorized taxonomic groups, such as viruses. The robustness and high accuracy observed for genome binning programs in the absence of strain diversity supported their application to large-scale data from various environments, recovering thousands of metagenome-assembled genomes^15–17^ (MAGs), and intensified efforts in advancing strain-resolved assembly and binning techniques. We here describe the results of the second round of CAMI challenges^18^, in which we assessed program performances and progress on even larger and more complex datasets, additionally including long-read data and assessment of key performance metrics such as runtime and memory use.

## Results

We created three comprehensive metagenome benchmark datasets representing a marine environment, a plant-associated environment that included fungal genomes and host plant material, as well as a very high strain diversity environment (“strain madness”). Datasets included both long and short-read data and were sampled from 1,680 microbial genomes and 599 circular elements of viruses and plasmids (Methods, Supplementary Table 1). Of these, 772 genomes and all circular elements were newly sequenced and distinct from taxa represented in public genome sequence collections (novel genomes), and the remainder were high-quality public genomes. Genomes were classified as “unique”, if they had an Average Nucleotide Identity (ANI) of less than 95% to any other genome, or “common”, if there were genomes with an ANI ≥ 95% in the benchmark data, as in the first CAMI challenge^6^. Overall, 901 genomes were unique: 474 for marine, 414 for plant-associated, and 13 for the strain madness data; and 779 were common: 303 for marine, 81 for plant-associated, and 395 for strain madness. In addition, a pathogen detection challenge was offered, based on a clinical metagenome sample from a critically ill patient with an unknown infection. Challenge participants were encouraged to submit reproducible results by providing executable software with exact parameter settings and reference databases used. Over all challenges, 5,002 results for 76 programs were received from 30 teams (Supplementary Table 2).

### Assembly challenge

Sequence assembly is a key component of metagenome analysis, with assemblies being subsequently used to recover genome and taxon bins. Assembly quality degrades for genomes with low evolutionary divergences, resulting in consensus or highly fragmented assemblies^19, 20^. Due to the relevance of strain-resolved assemblies for understanding microbial communities^21–23^, we assessed methods’ abilities to assemble strain-resolved genomes, together with the value of long and short-read data for assembly (Methods).

### Overall trends

We evaluated 155 submissions (Supplementary Table 2) for 20 assembler versions: A-STAR workflow (hybrid, contigs, and scaffolds), ABySS^24^ (short read, v.2.1.5), (meta)Flye^25^ (long read, v.2.4.1, v.2.8, v.2.8.1), (Meta)HipMer^26–28^ (short read, v.1.0, v.1.2.2, v.2.0, Metagenome, cgraph, cgraph-ono), GATB^29, 30^ workflow (hybrid, v.1.0), MEGAHIT^31^ (short read, v.1.1.2, v.1.1.4-2, v.1.2.7), Metahit_LINKS, Atlas^32^ (short read/hybrid single samples, v.2.1.0), (meta)SPAdes^33, 34^ (short read/hybrid, v.3.13.0, v.3.13.1, v.3.14-dev), OPERA-MS^35^ (hybrid, v.0.8.3, v.0.9), and Ray Meta^36^ (short read, v.2.3.1), including some with multiple settings and different data preprocessing options.

In addition, we created gold standard co- and single sample assemblies as in^19^, which included all regions covered by at least one read in the community-specific genome collections (i.e., marine, strain madness, plant-associated). The three gold standards of short, long, and hybrid marine data comprise 2.59 Gb, 2.60 Gb, and 2.79 Gb of assembled sequences, respectively, while the strain madness short, long, and hybrid gold standards consist of 1.45 Gb each.

Assemblies were evaluated with MetaQUAST v.5.1.0rc^37^, which was adapted for the evaluation of strain-resolved assembly (Supplementary text). To test the ability of the assemblers to generate near-complete strain-resolved genomes, we determined strain recall and precision, similar to^38^ (Supplementary Table 3). Strain recall measures how many genomes are recovered with high genome fraction and few mismatches (mm). Complementary to recall, strain precision assesses how accurately reference genomes are recovered, based on the fraction of correctly assembled high-quality, near-complete genomes (>90% genome fraction, <0.1% mm) divided by the overall number of assembled, near-complete genomes (>90% genome fraction). To facilitate comparisons, we ranked assemblies produced with different versions and parameter settings for a particular method based on key metrics (Methods) and chose the highest-ranking assembly as the representative (Fig. 1).

**Fig. 1:**
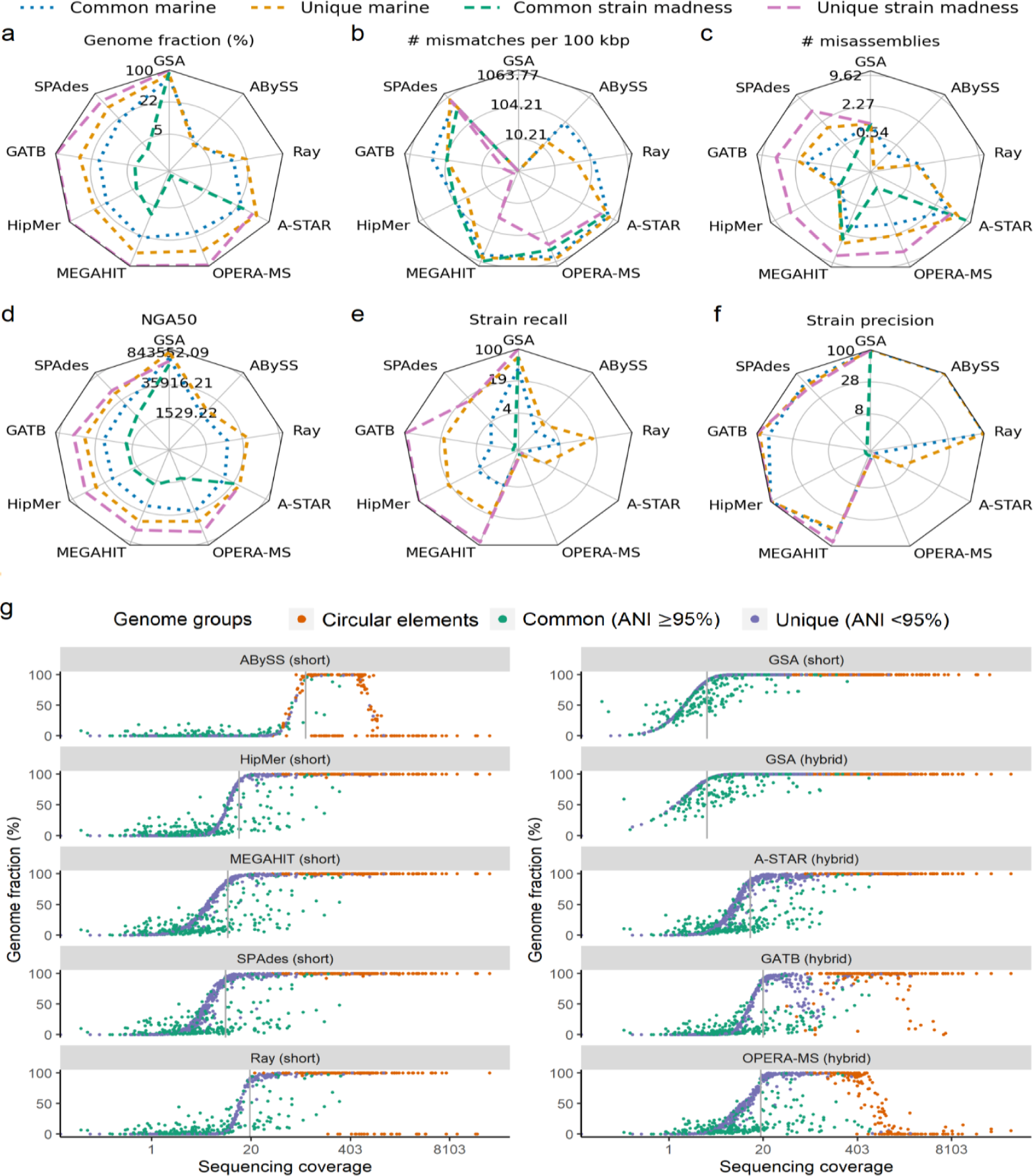
Radar plots of genome fraction (**a**), mismatches per 100 kb (**b**), misassemblies (**c**), NGA50 (**d**), strain recall (**e**), and precision (**f**) for assemblers on marine and strain madness data. For methods with multiple evaluated versions, the best ranked version on the marine data is shown. Absolute values for metrics are log scaled. Lines indicate different subsets of genomes analyzed, and the value of the gold standard assemblies (GSA) indicates the upper bound for a metric. The metrics are shown for both unique and common strain genomes. **g**, Genome recovery fraction versus genome sequencing depth (coverage) on the marine dataset. Blue indicates unique genomes (<95% ANI), green common genomes (ANI ≥95%), and orange high-copy circular elements. Grey lines indicate the coverage at which the first genome is recovered with ≥90% genome fraction.

Of these, short-read assemblers achieved genome fractions of up to 10.4% on the complete strain madness and 41.1% on marine data, both by MEGAHIT. The gold standard reported 90.8% and 76.9%, respectively (Fig. 1a, Supplementary Table 3). A- STAR excelled in terms of genome fraction on both data sets, but created more misassemblies and mm than others. HipMer had the fewest mm per 100 kb on the marine data set with 96 and GATB on the strain madness with 98 (Fig. 1b). The best hybrid assembler, A-STAR, improved the genome fraction to 44.1% on the marine dataset, at a cost of 773 mm/100 kb. The fewest mismatches (173) for hybrid assemblers were introduced by GATB. ABySS created the fewest misassemblies for the marine and GATB for the strain madness data (Fig. 1c). The most contiguous assemblies were provided by the hybrid assembler OPERA-MS for the marine data (Fig. 1d), with an average NGA50 of 28,244 across genomes, in comparison to 682,777 for the gold standard. The SPAdes hybrid submission had a higher NGA50 of 43,014, but it was not the best scoring SPAdes submission. A-STAR had the highest contiguity for the strain madness data (13,008 vs. 155,979 for gold standard). For short-read assembly, MEGAHIT had the highest contiguity on the marine (NGA50 of 26,599) and on the strain madness data (NGA50 of 4,793). Notably, differently from what we observed for the plant-associated long-read data (Supplementary Fig. 1), Flye performed less well than other assemblers across most metrics on the marine data, likely due to use of different versions or parameter settings (Supplementary Table 2).

HipMer ranked first across metrics on the marine data, as it produced few mm combined with a comparably high genome fraction and NGA50. On the strain madness data, GATB ranked best, with HipMer in second place. On the plant-associated dataset, HipMer again performed best across metrics, followed by Flye, whereby both outperformed the other short-read assemblers on this dataset in most metrics (Supplementary Fig. 1).

For several assemblers, preprocessing the data had a substantial impact on assembly quality (Supplementary Tables 2 and 3). In particular, using read quality trimming or error correction software, such as trimmomatic^39^, DUK^40^, Fastp^41^, or Bayeshammer^42^, improved assembly quality. Genome coverage was also a key factor for assembly quality (Fig. 1g). While both gold standards for short and hybrid assemblies contained genomes with more than 90% genome fraction starting at a coverage of 3.3x, the best assembler for low coverage genomes on the marine data set was SPAdes, recovering unique genomes starting at 9.2x. MEGAHIT, A-STAR, and HipMer required 10x, 13.2x, and 13.9x coverage, while Ray Meta recovered almost complete genomes from a coverage of 19.5x. Several assemblers reconstructed high copy circular elements well, with HipMer, MEGAHIT, SPAdes, and A-STAR reconstructing all of them (Fig. 1g). In comparison to well-performing software assessed in the first CAMI challenge, A-STAR substantially improved in genome fraction for the strain madness data by 20% (almost 3 times the genome fraction in relative terms) relative to MEGAHIT. HipMer introduced the fewest mismatches on the marine (67mm/100 kb) data. This was 30% less than Ray Meta, the best performing method also participating in CAMI 1. OPERA-MS improved on MEGAHIT in terms of NGA50 by 1,645 (6%), though using twice as much (long and short-read) data. SPAdes, which was not assessed in the first challenge, consistently was among the top submissions for most metrics.

### Closely related genomes

The first CAMI challenge revealed substantial differences in assembly quality between unique and common strain genomes^6^. We further focused on this by providing a dataset consisting almost entirely of common strain genomes. The upper bound for strain recall was 54.9%, as provided by the marine gold standard assembly, and 67.4% for the strain madness one (Fig. 1e). Strain recall varied little across several evaluated threshold settings for genome fraction (>90%, >75%) and mismatches (<0.1% mm/kb; <0.5% mm/kb). Therefore, we set >90% genome fraction and <0.1% mm/kb as thresholds.

Overall, GATB ranked best across metrics on strain madness data and the common strain madness genomes, while HipMer ranked best on marine data and common marine genomes (Supplementary Table 3). HipMer had the highest strain recall (14.4% on marine, 3.2% on strain madness), similar or even better than the best hybrid assembler, GATB (10.8% on marine, 2.9% on strain madness). Multiple assemblers - ABySS, HipMer, and Ray Meta on marine and GATB on strain madness - achieved 100% precision (Fig. 1f). For strain madness common genomes, A-STAR recovered the most, with 1.5% recall (<0.5% mm, >75% genome fraction) and 23.1% strain precision. HipMer recovered a lower genome fraction (4.1% versus 30.4% for A-STAR), but also created fewer misassemblies per genome (0.5) and mismatches (0.1%), resolving fewer, but higher quality strain genomes with high precision (0.8% strain recall, 100% strain precision). A major difference between common genomes in the two datasets is that, in the strain madness data, virtually all genomes form a single cluster with >95% ANI, while the marine data includes multiple clusters with a few closely related genomes each. Accordingly, strain recall was higher for common marine than common strain madness genomes (Supplementary Table 4). On marine common genomes, SPAdes had the highest strain recall (8.7%) and 96.7% strain precision, followed by A-STAR (7.5% recall, 69.4% precision). A-STAR (26.7%) and MEGAHIT (24.6%) achieved the highest genome fractions.

Across metrics, unique genome assemblies were superior to common genome ones (Fig. 1, Supplementary Table 4), for marine genomes by 15.7% in strain recall, 22.7% genome fraction, 3-fold NGA50, except in strain precision (-1.2%), on average (Supplementary Table 5), resulting in substantially more complete, higher quality, and less fragmented assemblies. A-STAR provided the most complete assemblies (55.3% genome fraction), the HipMer assembly had the highest strain recall (20.4%), HipMer, ABySS, Flye, and Ray Meta the highest strain precision (100%) and OPERA-MS an exceptional average NGA50 (187,083, 75% of the gold standard NGA50). HipMer ranked best across all metrics, with 100% strain recall and precision, 98.5% genome fraction, and the fewest mismatches (0.001%). Unique strain madness genomes were recovered best, with on average 81.4% strain recall, 89.4% strain precision, 86.4% genome fraction, and an NGA50 of 120,771 (Supplementary Table 6).

### Difficult to assemble regions

As the marine data also include high quality public genomes, assembly performance can be assessed for particularly difficult to assemble genomic regions, such as repeats or highly conserved elements (e.g., 16S). To assess recovery of complex regions, we selected 50 unique, public genomes present as a single contig in the gold standard and with annotated 16S sequences. We mapped assembly submissions to these 16S sequences and measured their completeness and gap-compressed divergence (Supplementary Fig. 2). A-STAR partially recovered 102 (78%) of 131 16S gold standard sequences. The hybrid assemblers GATB (mean recovered gene fraction 60.1%) and OPERA-MS (mean 47.1%) recovered the most complete 16S sequences. The mean fraction of genes recovered by short-read assemblers spanned between 29.6% (HipMer) and 36.9% (MEGAHIT) and were very accurate for ABySS and HipMer (<1% divergence). Average assembly quality was better for public than for novel genomes in key metrics, such as genome fraction and NGA50 (Supplementary Fig. 3).

### Single versus co-assembly

For multi-sample metagenome datasets, there are two assembly strategies: pooling samples (coassembly) and single-sample assembly^16, 28, 43^. We evaluated the assembly quality of both strategies for genomes that were spiked into the plant-associated data with specific coverages (Supplementary Table 8) across submitted results for five assemblers (Supplementary Fig. 4). Two genomes were unique with 8x coverage across the pooled samples, distributed into 16 samples or one sample, respectively. For the genome split across 16 samples, only HipMer recovered it from the pooled samples, while the genome present in only one sample was reconstructed well by all assemblers from single and pooled samples. For genomes unique to a single sample, but common in pooled samples (LjRoot109, LjRoot170), HipMer showed a better performance on the single samples, while OPERA-MS generally performed better on the pooled samples (Supplementary Fig. 4), and other assemblers traded a higher genome fraction against more mismatches. Thus, co-assembly could improve assembly for OPERA-MS in general and for short-read assemblers on low coverage genomes without expected strain diversity across samples. For HipMer, single-sample assembly might be preferable, if coverage is sufficient and closely related strains are expected.

### Genome binning challenge

Genome binners group contigs or reads together to recover genome bins from metagenome data. We evaluated 95 results for 18 binning software versions on short read-assemblies: 22 for the strain madness gold standard assemblies (GSA), 17 for the strain madness MEGAHIT assembly (MA), 19 for marine MA, 15 for the marine GSA, 12 for the plant-associated GSA, and 10 for the plant-associated MA (Supplementary Tables 9-15, Supplementary Table 2 for software availability, versions, and parameters). In addition, 7 results on the plant-associated hybrid assemblies were evaluated. Methods included well performing ones from the first CAMI challenge and popular software, such as MetaBAT^44, 45^ (v.2.15-5, v.2.13-33, v.0.25.4), MaxBin^46^ (v.2.2.7, v.2.0.2) and CONCOCT^47^ (v.1.1.0, v.0.4.1), as well as Autometa^48^ (v.cami2), LSHVec^49^ (v.cami2), MetaBinner (v.1.3, v.1.2, v.1.1, v.1.0), UltraBinner (v.1.0), MetaWRAP^50^ (v.1.2.3), SolidBin^51^ (v.1.3), and Vamb^52^ (v.3.0.1, v.fa045c0). While for GSA contigs, the ground truth genome assignment is known, for the MA, we considered the ground truth for a contig to be the best matching genomes identified using MetaQUAST v5.0.2 with default settings. We assessed the average purity of bins and completeness of genomes (and their summary using the F1-score), the number of high-quality genomes recovered, as well as the Adjusted Rand Index (ARI) for the binned data, using AMBER v.2.0.3^53^ (Methods). The ARI, in combination with the fraction of binned data, quantifies binning performance for the overall dataset.

The performance of genome binners varied widely across metrics, software versions, datasets, and assembly type (Fig. 2), while parameters affected performance mostly by less than 3%. For the marine GSA, average bin purity was 81.3±2.3% and genome completeness was 36.9±4.0% (Fig. 2a,b, Supplementary Table 9). For the marine MA, average bin purity (78.3±2.6%) was similar, while average completeness was only 21.2±1.6% (Fig. 2a,c, Supplementary Table 10), due to many short contigs with 1.5-2 kb, which most binners did not attempt to bin (Supplementary Fig. 5). For the strain madness GSA, average purity and completeness decreased, by 20.1% to 61.2±2.3%, and by 18.7% to 18.2±2.2%, respectively, relative to the marine GSA (Fig. 2a,d, Supplementary Table 11). While the average purity on the strain madness MA (65.3±4.0%) and GSA were similar, the average completeness dropped further to 5.2±0.6%, again due to a larger fraction of short contigs not binned (Fig. 2a,e, Supplementary Table 12). For the plant-associated GSA, purity was almost as high as for marine (78.2%±4.5; Fig. 2a,f, Supplementary Table 13), but bin completeness decreased relative to other GSAs (13.9±1.4%), due to poor recovery of the mostly low abundant, large, fungal genomes. Notably, the *A. thaliana* host genome (5.6x coverage) as well as fungi with more than 8x coverage were binned with much higher completeness and purity than genomes with lower coverage (Supplementary Fig. 6). Binning of the hybrid assembly further increased average purity to 85.1±6.3%, while completeness remained similar (11.9±2.1%, Supplementary Table 14). For the plant-associated MA, average purity (83±3.3%) and completeness (12.4±1.5%; Fig. 2a,g, Supplementary Table 15) were similar to the GSA.

**Fig. 2:**
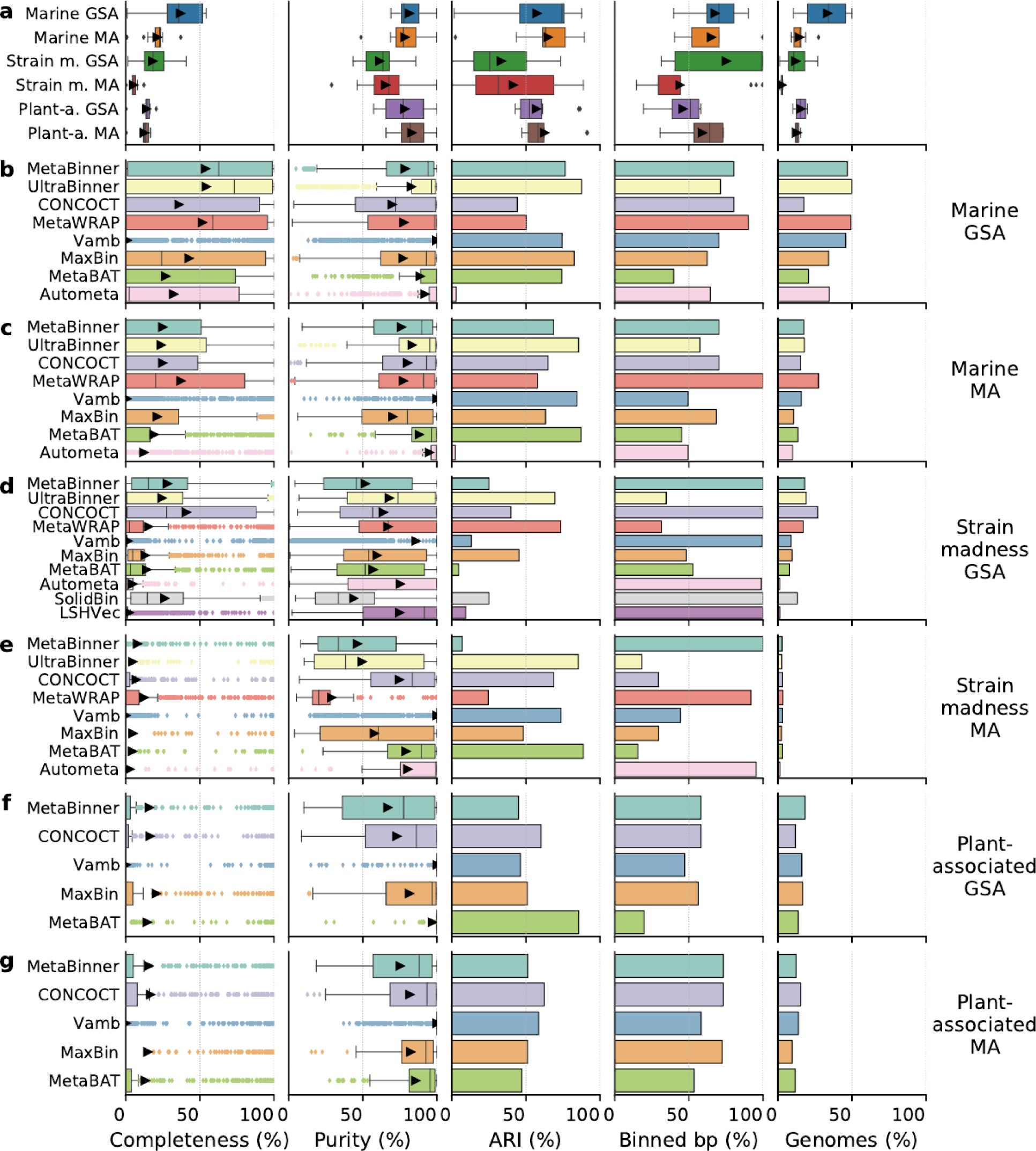
Performance of genome binners on short-read assemblies (GSA: gold standard, MA: MEGAHIT) of the marine, strain madness, and plant-associated data. a, Boxplots of average completeness, purity, Adjusted Rand Index (ARI), percentage of binned bp, and fraction of genomes recovered with moderate or higher quality (>50% completeness, <10% contamination) across methods from each dataset (Methods). Arrows indicate the average. **b-g,** Boxplots of completeness per genome and purity per bin, and bar charts of ARI, binned bp, and moderate or higher quality genomes recovered, by method, for each dataset. The submission with the highest F1-score per method on a dataset is shown (Supplementary Tables 9-15).

To quantitatively assess binners across gold standard and real assemblies for the three datasets, we ranked the best submissions (Supplementary Tables 16-19) across metrics (Methods). For marine and strain madness, the best trade-off performances were given by CONCOCT and MetaBinner for MAs, UltraBinner for GSAs, and MetaBinner overall. CONCOCT also performed best on plant-associated assemblies. UltraBinner had the best completeness on the marine GSA, CONCOCT on the strain madness GSA and plant-associated MA, MetaWRAP on marine and strain madness MAs, and MaxBin on the plant-associated GSA. Vamb had the best purity in all settings, while UltraBinner had the best ARI for the marine GSA, MetaWRAP for the strain madness GSA, and MetaBAT for MAs and all plant-associated assemblies. MetaWRAP and MetaBinner assigned the most for the marine and plant-associated assemblies, respectively, and many methods assigned all strain madness contigs, though with low ARI (Fig. 2b-g). UltraBinner recovered the most high-quality genomes from the marine GSA, MetaWRAP from the marine MA, CONCOCT from strain madness assemblies and plant-associated GSA, and MetaBinner from the plant-associated GSA and hybrid assemblies (Fig. 2, Supplementary Table 20). For plasmids and other high copy circular elements, Vamb performed best, with an F1-score of 70.8%, 54.8% completeness, and 100% purity, while the next best method, MetaWRAP, had an F1-score of 12.7% (Supplementary Table 21).

### Effect of strain diversity

For both marine and strain madness GSAs, binning of unique strains was substantially better than for common strains (Supplementary Fig. 7, Supplementary Tables 9 and 11). Differences were more pronounced on strain madness, for which unique strain bin purity was particularly high (97.9±0.4%). The best ranking across metrics and these four data partitions was obtained by UltraBinner for unique genomes and overall, as well as CONCOCT for common strains (Supplementary Table 22). UltraBinner had the highest completeness on unique strains, while CONCOCT ranked best for common strains and across all partitions together. Vamb ranked first by purity in all settings, UltraBinner by ARI, and MetaBinner by most assigned. Due to the dominance of unique strains in the marine and common strains in the strain madness dataset, the best binners in the respective data were the same as for the entire datasets (Supplementary Tables 9 and 11) and performances similar for most metrics.

### Taxonomic binning challenge

Taxonomic binners group sequences into bins labelled with a taxonomic identifier. For taxonomic binning, we evaluated 547 results for nine methods and versions: LSHVec v.cami2^49^, PhyloPythiaS+ v.1.4^54^, Kraken v.2.0.8-beta^55^ and v.0.10.5-beta (*cami1*), DIAMOND v.0.9.28^56^, MEGAN v.6.15.2^57^, Ganon v.0.1.4 and v.0.3.1^58^, and NBC++^59^. Of these, 75 were for the marine, 405 for strain madness, and 67 for plant-associated data, on either reads or gold standard assemblies (Supplementary Tables 2). We assessed the average purity and completeness of bins and the accuracy per sample at different taxonomic ranks (Methods).

On the marine data, average taxon bin completeness across ranks was 63%, average purity 40.3%, and accuracy per sample bp 74.9% (Fig. 3a, Supplementary Table 23). On the strain madness data, accuracy was similar (76.9%; Fig. 3b, Supplementary Table 24), while completeness was ∼10% higher and purity lower by that much. On the plant-associated data, purity was between those of the first two datasets (35.%), but completeness and accuracy were lower (44.2% and 50.8%, respectively; Fig. 3c, Supplementary Table 25). For all datasets, performances declined at lower taxonomic ranks, most notably from genus to species rank by 22.2% in completeness, 9.7% in purity, and 18.5% in accuracy, on average.

**Fig. 3:**
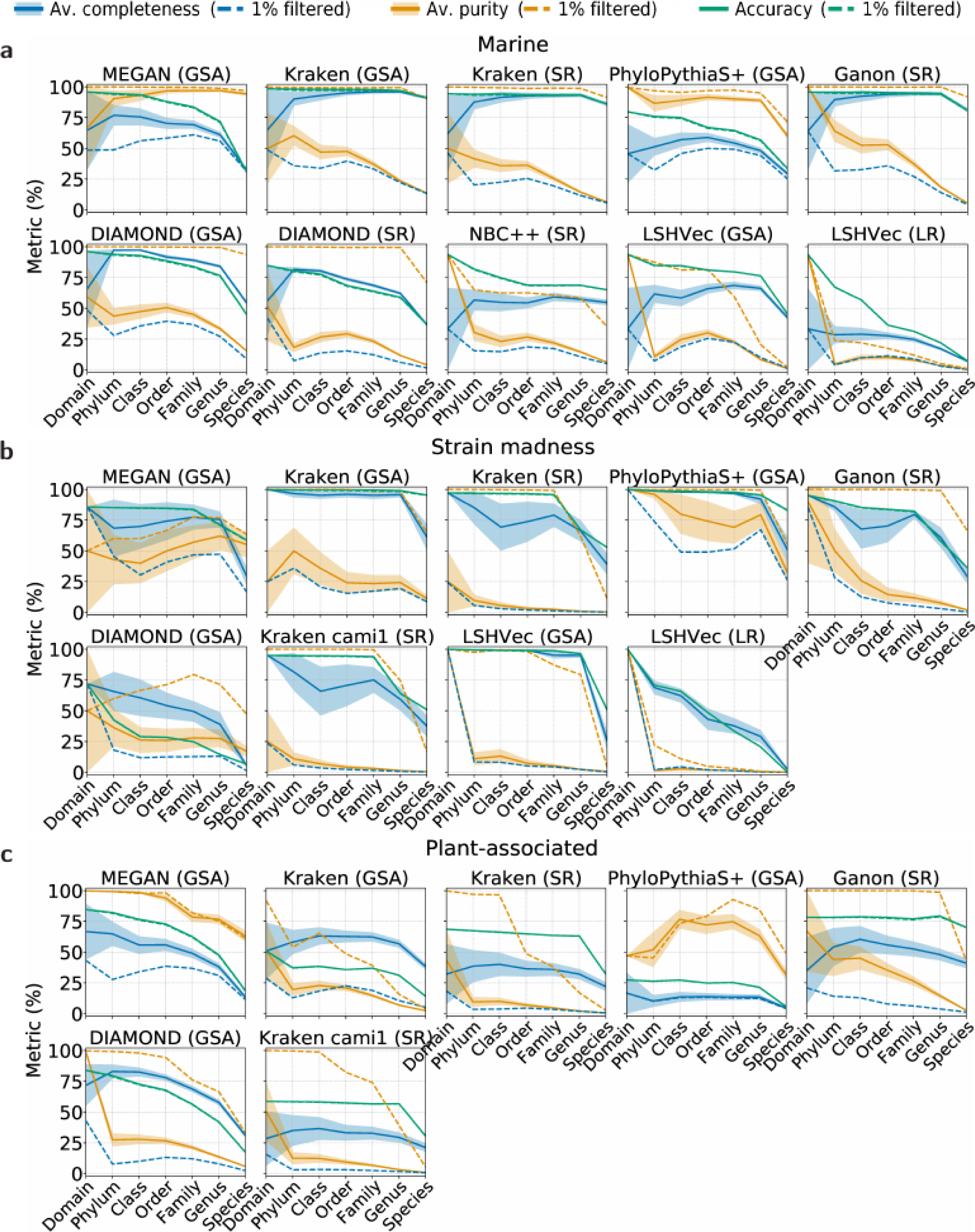
Taxonomic binning performance across ranks for the marine (a), strain madness (b), and plant-associated datasets (c). Metrics were computed over unfiltered (solid lines) and 1%-filtered (i.e., without the 1% smallest bins in bp, dashed lines) predicted bins of short reads (SR), long reads (LR), and contigs of the gold standard assembly (GSA). Shaded bands show the standard error across bins.

Across all datasets, MEGAN on contigs ranked first across metrics and all ranks (Supplementary Table 26), closely followed by Kraken v.2.0.8-beta on contigs and then by Ganon on short reads. Kraken v.2.0.8-beta on contigs was the best for genus and species, and on marine data in completeness and accuracy (89.4%, 96.9%; Supplementary Table 23 and 27). Due to the presence of public genomes, Kraken’s completeness on marine data was much higher than in the first challenge, particularly at species and genus rank (average of 84.6% and 91.5%, respectively, compared to 50% and 5%), while purity remained similar. MEGAN on contigs ranked highest for taxon bin purity on the marine and plant-associated data (90.7%, 87.1%; Supplementary Tables 23, 25, 27, and 28). PhyloPythiaS+ ranked best for the strain madness data across metrics, as well as in completeness (90.5%) and purity (75.8%) across ranks (Supplementary Tables 24 and 29). DIAMOND on contigs ranked best for completeness (67.6%) and Ganon on short reads for accuracy (77.1%) on the plant-associated data.

Filtering of the 1% smallest predicted bins per taxonomic level is a popular post-processing approach. On all datasets, filtering increased average purity to above 71% and reduced completeness, to ∼24% on marine and strain madness and 13.4% on plant-associated data (Supplementary Table 23-25). Accuracy was not much affected, as large bins contribute more to this metric. Kraken on contigs still ranked first in filtered accuracy and MEGAN across all filtered metrics (Supplementary Table 26). MEGAN on contigs and Ganon on short reads profited the most from filtering, ranking first in filtered completeness and purity, respectively, across all datasets and taxonomic levels.

### Taxonomic binning of divergent genomes

To investigate the effect of increasing divergence between query and reference sequences for reference-based taxonomic binners, we categorized genomes by their distances to public genomes (Supplementary Fig. 8, Supplementary Tables 30 and 31). Sequences of known marine strains were assigned particularly well at the species rank by Kraken (accuracy, completeness, and filtered purity above 93%) and MEGAN (91% purity, 33% completeness and accuracy). Kraken also best classified new strain sequences at species level, though with less completeness and accuracy for the marine data (68% and 80%, respectively). It also had the best accuracy and completeness across ranks, but low unfiltered purity. For the strain madness data, PhyloPythiaS+ performed similarly well up to genus level, and best assigned new species at genus level (93% accuracy and completeness, and 75% filtered purity). Only DIAMOND classified viral contigs, though with low purity (50%) and completeness and accuracy (both 3%), and no method classified sequences as plasmids.

### Taxonomic profiling challenge

Taxonomic profilers quantify the presence and relative abundances of microbial community taxa from metagenome samples. This is in contrast to taxonomic sequence classification, which assigns taxon labels to individual sequences and results in taxon-specific sequence bins (and sequence abundance profiles), instead of taxonomic abundance profiles for entire samples or datasets^60^. We evaluated 4,195 profiling results (292 marine, 2,603 strain madness, and 1,300 plant-associated datasets), from 22 method versions (Supplementary Table 2) with the majority of the results originating from short-read samples, and a few from long-read samples, assemblies, or averages across samples. Performance was evaluated with OPAL v.1.0.10^61^ (Methods). The quality of predicted taxon profiles was determined based on completeness and purity of identified taxa, relative to the underlying ground truth, for individual ranks, while taxon abundance estimates were assessed using the L1 norm error for individual ranks, and the weighted Unifrac error across ranks. To assess alpha diversity of profiling results, the absolute difference between predicted and actual Shannon equitability index was determined (Methods).

### Taxon identification

On the marine data, methods performed well until genus rank (average purity 70.4% and completeness 63.3%), with a substantial drop at species level, to 44.4% purity and 47.1% completeness (Supplementary Fig. 9, Supplementary Tables 32 and 33). mOTUs v.2.5.1^62^ had completeness and purity above 80% at genus and species ranks, and Centrifuge^63^ and MetaPhlAn 2.9.22^64, 65^ just at the genus rank (Fig. 4). Other methods with completeness above 80% at either rank were Bracken^66^, Centrifuge v.1.0.4 beta, MetaPhlAn v.2.9.22, and NBC++^59^, while CCMetagen^67^, DUDes 0.08^68^, LSHVec gsa^49^, Metalign^69^, MetaPalette^70^, and MetaPhlAn cami1 had more than 80% purity. Filtering out the rarest (1%) predicted taxa per rank decreased completeness by ∼22%, while increasing precision by ∼11%.

**Fig. 4:**
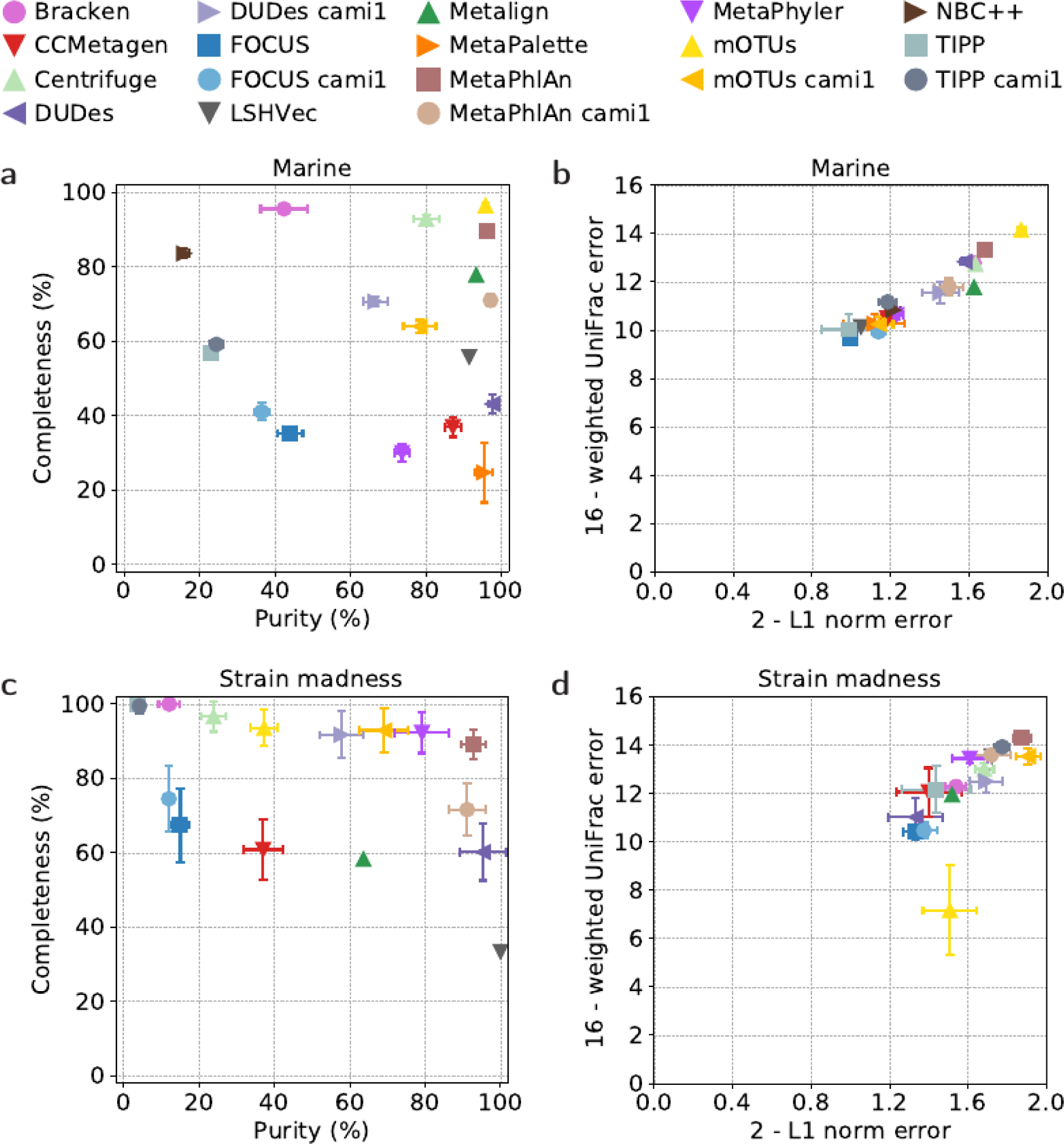
Taxonomic profiling results for the marine (a, b) and strain madness (c, d) datasets at genus level. Results are shown for the overall best ranked submission per software version (Supplementary Tables 33 and 35). **a, c,** purity vs. completeness. **b, d,** upper bound of L1 norm minus actual L1 norm vs. upper bound of weighted UniFrac error minus actual weighted UniFrac error. Error bars show the standard deviation across samples. Metrics were determined using OPAL with default settings.

A similar trend was evident for the strain madness data, with methods performing well until the genus rank (average purity 52.1% and completeness 80.5%), and a substantial drop at species rank (average purity 22.3%, completeness 43.9%, Supplementary Tables 34 and 35). In comparison to the marine data, the average purity per method across ranks fell from 75.3% to 57.1%, while completeness rose from 72% to 79%. At the genus rank, particularly MetaPhlAn v.2.9.2 (89.2% completeness, 92.8% purity), MetaPhyler v.1.25^71^ (92.3% completeness, 79.2% purity) and mOTUs of the first CAMI challenge (92.9% completeness, 69.1% purity) performed well, but no method excelled at the species rank. DUDes v.0.08 and LSHVec gsa also had high purity, while Centrifuge v.1.0.4 beta, DUDes v.cami1, and TIPP v.4.3.10 and v.cami1 had high completeness.

Also for the plant-associated data, average purity and completeness decreased considerably from genus to species, from 62.9% to 31.7% and from 42.1% to 31.9%, respectively (Supplementary Tables 36 and 37). The average purity across ranks was the highest among the datasets (77.8%), while average completeness was the lowest (49.7%), due to the many newly sequenced bacterial and fungal genomes (Methods), which most methods failed to detect. Bracken v.2.6 performed best for completeness across ranks (average 63.7%) and “sourmash gather 3.3.2 k21”^72^ on short-reads for species (53.8%). “Sourmash gather 3.3.2 k21” on Pacific Biosciences reads and MetaPhlAn v.3.0.7 on short reads had the highest purity across ranks (94.7%, 94.67%), and the latter also for species (68.8%).

### Relative abundances

Abundances across ranks and submissions were on average predicted better for strain madness than marine data, which has less complexity above the strain level, with the L1 norm error improving from 0.44 to 0.3, and average weighted UniFrac error from 4.65 to 3.79 (Supplementary Table 32, 34, 36). Abundance predictions were not as good on the plant-associated data and averaged 0.57 in L1 norm error and 5.16 in weighted UniFrac. On the marine data, mOTUs v.2.5.1 had the lowest L1 norm error at almost all levels and 0.12 on average, including the lowest at genus and species with 0.13 and 0.34, respectively. It was followed by MetaPhlAn v.2.9.22, with 0.22 on average, and 0.32 and 0.39 at genus and species. Both methods also had the lowest weighted UniFrac error, followed by DUDEs v.0.08. On the strain madness data, mOTUs cami1 performed best in L1 norm error across ranks with 0.05, and also at genus and species with 0.1 and 0.15, followed by MetaPhlAn v.2.9.22 with 0.09 on average, and 0.12 and 0.23 at genus and species. The latter also had the lowest weighted UniFrac error, followed by TIPP v.cami1 and mOTUs v.2.0.1. On the plant-associated data, Bracken v.2.6 had the lowest L1 norm error across ranks, with 0.36 on average, and at genus, with 0.34. “sourmash gather 3.3.2 k31” on short-reads had the lowest at species, with 0.55. Both methods also had the lowest UniFrac error on this dataset. Several methods also accurately reconstructed the alpha diversity of samples using the Shannon equitability; best (0.03 or less absolute difference to gold standards) across ranks were mOTUs v.2.5.1, DUDes v.0.08, MetaPhlAn v.2.9.22, as well as the versions of the first CAMI challenge of DUDes v.0.08 and MetaPhlAn on marine data, together with DUDes v.cami1 and MetaPhlAn v.2.9.22 on strain madness data. On the plant-associated data, mOTUs v.cami1 and Bracken v.2.6 performed best with this metric (0.08 and 0.09).

### Difficult and easy taxa

For all methods, viruses, plasmids, and Archaea were particularly difficult to detect (Supplementary Fig. 10, Supplementary Table 38) in the marine data. While many Archaeal taxa were detected by several methods, some taxa, such as Candidatus Nanohaloarchaeota, were not detected by any method in any sample. Similarly, no method detected any plasmids or viruses. In contrast, bacterial taxa in the Terrabacteria group and the phyla of Bacteroidetes and Proteobacteria were correctly detected by each method in all samples.

### Method similarity

To assess software performances in relation to their methodological similarity, we clustered submissions based on the Bray Curtis dissimilarity on the vectors of precision and recall per taxa, averaged over ranks. Methods using similar information types, e.g., k-mer based (NBC++, Bracken), alignment (CCMetagen, Metalign), and marker gene approaches (mOTUs, MetaPhlAn) tended to cluster (Supplementary Fig. 11); for example, the two alignment-based approaches are more similar to each other than to other methods. Interestingly, the marker gene approaches are most similar to the gold standard, suggesting this class of methods is particularly well suited to infer taxonomic profiles.

### Clinical pathogen prediction challenge

A short-read metagenomic sequencing dataset of a blood sample from a patient with hemorrhagic fever of unknown cause was provided for participants to identify a causal pathogen together with further pathogens. Ten manually curated, hence not fully reproducible results were received (Supplementary Table 39). The total number of identified taxa per result varied considerably (Supplementary Fig. 12). Three submissions correctly identified the causal pathogen, Crimean-Congo hemorrhagic fever orthonairovirus (NCBI taxid 1980519), using the taxonomic profilers MetaPhlAn v.2.2, Bracken v.2.5, and CCMetagen v.1.1.3^67^. Another submission using Bracken v.2.2 correctly identified orthonairovirus, but without indicating it as the causal pathogen.

### Computational requirements

We measured the runtimes and maximum main memory usage for submitted methods across the marine and strain madness data (Fig. 5, Supplementary Table 40, Methods). Compute and memory efficient methods capable of processing the entire datasets within minutes to a few hours were available in every method category, even including some of the identified top ranked techniques. Substantial differences were seen within categories and even between versions, ranging from methods executable on standard desktop machines to those requiring extensive hardware and heavy parallelization. Of the assemblers, MEGAHIT was the fastest and most memory efficient, requiring 7 h and 42 GB of main memory to process marine short reads. This was 30% and ∼25% less time and memory than required by the second fastest and most memory efficient methods, OPERA-MS and GATB, respectively. On the marine assemblies, genome binners on average required ∼3x less time than for the smaller strain madness assemblies (29.2 h vs. 86.1 h), but used almost 4x more memory (69.9 GB vs. 18.5 GB). MetaBAT 2.13.33 was the fastest (1.07 and 0.05 h) and most memory efficient genome binner (max. memory usage 2.66 GB, 1.5 GB) on both datasets. It was ∼5x and ∼635x faster than the second fastest method, Vamb fa045c0, ∼6x faster than LSHVec 1dfe822 on marine, and 765x faster than SolidBin 1.3 on strain madness data; ∼2x and ∼5x more memory efficient than next ranking MaxBin 2.0.2 and CONCOCT 1.1.0 on marine data, respectively. Both MetaBAT and CONCOCT were substantially (∼11x and ∼4x) faster than the versions assessed in CAMI 1. Like genome binners, taxonomic binners ran longer on the marine than the strain madness assemblies, e.g., PhyloPythiaS+ with 287.3 vs. 36 h, respectively, but had a similar or slightly higher memory usage. On the marine raw read data, taxon profilers, on the other hand, were almost 4x faster on average (16.1 h vs. 60.8 h) than on the 10x larger strain madness read dataset, but used more memory (38.1 GB vs. 25 GB). The fastest and most memory efficient taxonomic binner was Kraken, requiring only 0.05 and 0.02 h, respectively, and ∼37 GB memory on both datasets, for reads or contigs. It was followed by DIAMOND, which ran ∼500x and ∼910x as long on the marine and strain madness gold standard assemblies, respectively. FOCUS 1.5 and Bracken 2.2 were the fastest profilers on the marine (0.51 and 0.66 h, respectively) and strain madness (1.89, 3.45 h) data. FOCUS 1.5 also required the least memory (0.16 GB for marine, 0.17 GB for strain madness), followed by mOTUs 1.1.1 and MetaPhlAn 2.2.0.

**Fig. 5:**
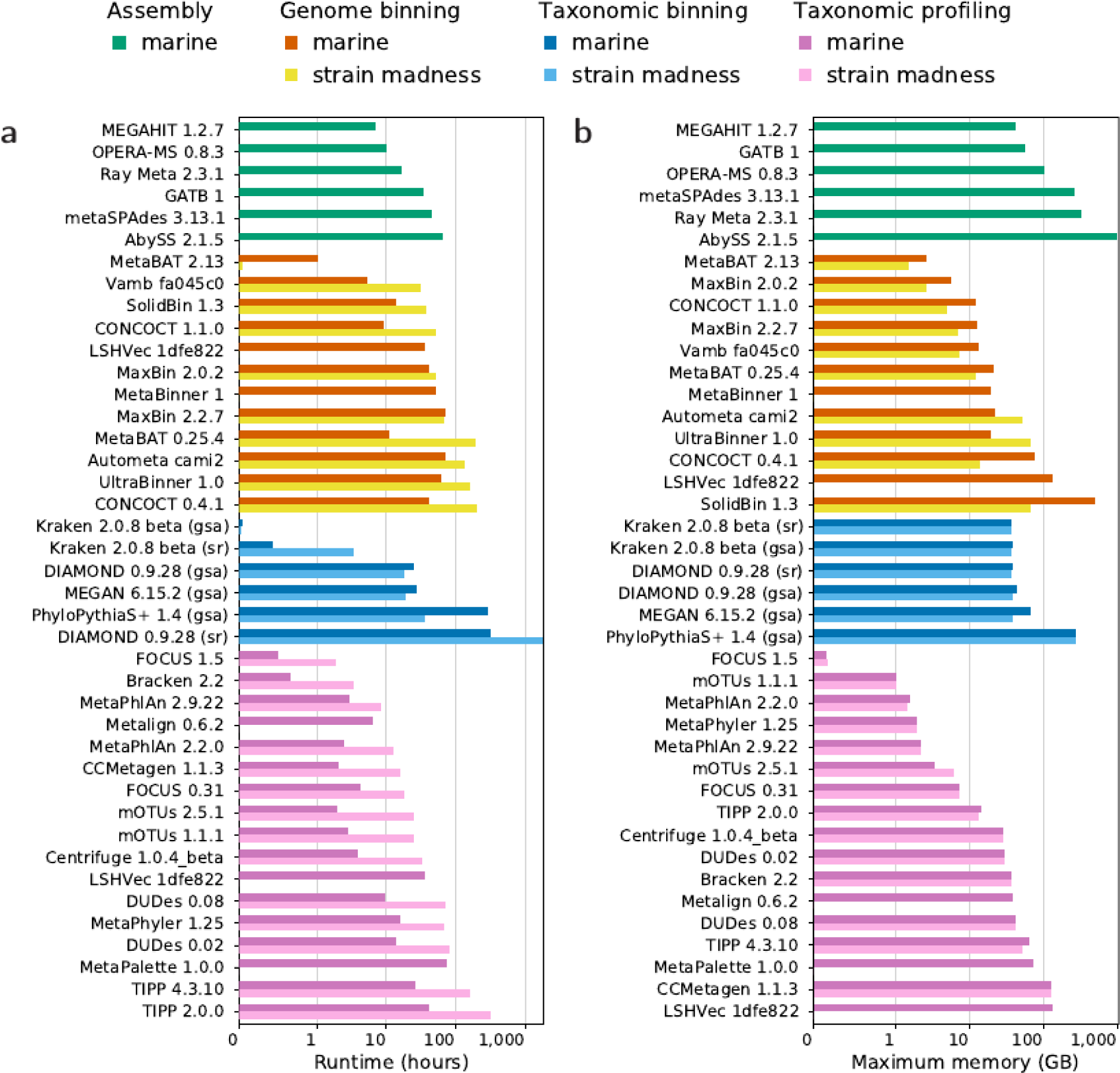
Runtime (**a**) and maximum memory usage (**b**) of software from all categories for the marine and strain madness data (Supplementary Table 40) with log-scaled x-axis.

## Discussion and conclusions

Assessing metagenomic analysis software thoroughly, comprehensively, and with little bias is key for optimizing data processing strategies as well as tackling open challenges in the field. In its second round, CAMI offered a diverse set of benchmarking challenges across a comprehensive data collection reflecting recent technical developments of the field. Here, we analyzed 5,002 results of 76 program versions with different parameter settings across 131 long and short read metagenome samples from four environments (marine, plant-associated, strain madness, clinical pathogen challenge). This effort increased the number of results 22x and the number of benchmarked software versions 3x in comparison to the first CAMI challenge, delivering extensive new insights into software performances and their interpretation across a wide range of conditions. By systematically assessing runtime and memory requirements, we added two more key performance dimensions to the benchmark, which are very important to consider, given the ever-increasing dataset sizes.

In comparison to software assessed in the first CAMI challenges, assembler performances rose for best performing methods by up to 20%. Still, in the presence of closely related strains, assembly contiguity, the fractions of genomes recovered, and strain recall decreased, suggesting that most assemblers, sometimes intentionally^31, 34^, did not accurately resolve strain variation, resulting in more fragmented, less strain-specific assemblies. In addition, genome coverage, parameter settings, and data preprocessing notably impacted assembly quality, while performances did not differ much across software versions. Most submitted metagenome assemblies used only short reads, and long and hybrid assemblies had no higher overall quality. Hybrid assemblies, however, were better for difficult to assemble regions, such as the 16S rRNA gene, recovering substantially more complete genes than most short-read submissions. Hybrid assemblers were also less affected by the presence of closely related strains in pooled samples, suggesting that the addition of long reads helps to distinguish individual strains.

In comparison to the first CAMI challenges, ensemble binners presented a development accompanied by substantial improvements across metrics in comparison to most individual methods. Overall, genome binners demonstrated variable performances across different metrics and dataset types, with both high strain diversity and assembly quality presenting challenges that substantially reduced performances relative to unique genome binning, even in the case of large sample numbers, such as for the strain madness dataset. Interestingly, for the plant-associated data, which also included plant host and 55 fungal genomes, given sufficient coverage, high-quality bins were obtained also for these taxa.

For taxonomic binners and profilers, highly performant and computationally efficient software was available, performing well across a range of conditions and metrics. Particularly the profiling field has matured in comparison to the first challenges, with less variance in top performing methods across taxon identification, abundance, and diversity estimates. Performance for both categories was found to be high for genus rank and above, with a substantial drop for bacterial species. As the second challenge data include high quality public genomes, the data is less divergent from publicly available data than for the first challenges, on which method performances declined already going from family to genus rank. It was also low for Archaea, viruses and plasmids, suggesting a need for developers to extend their focus in terms of reference sequence collections and model development. Another encouraging result is that the causal pathogen was successfully identified by several submissions in a clinical pathogen challenge. However, due to manual curation, none was reproducible, which indicates another area requiring improvement.

In its second challenge, CAMI identified key advances across common metagenomics software categories as well as current challenges for the field. As the state-of-the-art in methods and data generating techniques progresses, it will be important to continuously reevaluate these questions. In addition, computational methods for other microbiome data modalities^12^ and multi-omics data integration could be jointly assessed. Most importantly, CAMI is a community-driven effort, and we encourage everyone with an interest in benchmarking from the field of microbiome research to join us.

## Methods

### Community involvement

We gathered community input on the nature and principles of implementing benchmarking challenges and datasets in public workshops and hackathons (https://www.microbiome-cosi.org/cami/participate/schedule). The most relevant metrics for performance evaluation and data interpretation were discussed in a public evaluation workshop with challenge participants and developers of evaluation software, where first challenge results were presented in an anonymized manner. Computational support for challenge participants was provided by the de.NBI cloud.

### Standardization and reproducibility

To ensure reproducibility and assess computational behavior (runtimes and memory consumption) of the software used to create challenge submissions, we reproduced and reassessed the results according to submission specifications (Supplementary Table 2, https://data.cami-challenge.org/). For metagenome assemblers, computational requirements were assessed on a machine with Intel Xeon Processor (2.6GHz) virtualized to 56 cores (50 cores were used) and 2755 GB of main memory and for binners and profilers on a machine with an Intel Xeon E5-4650 v4 CPU (virtualized to 16 CPU cores, 1 thread per core) and 512 GB of main memory. We also updated Docker BioContainers implementing a range of commonly used performance metrics to include all metrics used in this evaluation (Supplementary Table 2).

### Genome sequencing and assembly

Illumina paired-end read data of 796 newly sequenced genomes, of which 224 stem from a *Arabidopsis thaliana* root environment, 176 from a marine environment^73^, 384 clinical *Streptococcus pneumoniae* strains, and twelve strains from a murine gut environment were assembled using a pipeline with the SPAdes^33^ metagenome assembler (version 3.12). We removed contigs smaller than 1 kb, as well as genome assemblies with a contamination of 5% or more and completeness of 90% or less, as determined with CheckM^74^ version 1.011. Newly assembled and database genomes were taxonomically classified with CAMITAX^75^ and used as input for microbial community and metagenome data simulation with CAMISIM^76^, based on the *from_profile* mode for the marine and plant-associated dataset and the *de novo* mode for the strain madness datasets. All scripts and parameters for these steps are provided in the Supplementary Material and on GitHub (https://github.com/CAMI-challenge/second_challenge_evaluation/tree/master/scripts/data_generation).

For the plasmid dataset, inlet wastewater from a wastewater treatment plant on Zealand, Denmark was used to generate a plasmid sample similar to the procedure in^77^. Sequencing was performed on a NextSeq 500 on Nextera sequencing libraries (Illumina, San Diego, California, USA). A bioinformatic workflow described in^78^ was used to identify complete circular plasmids above 1 kb in size in the dataset.

### Challenge datasets

For the challenges, participants were provided with long and short-read sequences for two metagenome datasets representing a marine and a plant associated environment, respectively, in complexity and taxonomic distribution, and for a “strain madness” dataset with very high strain diversity. Furthermore, a short-read clinical metagenomic dataset from a critically ill patient was provided.

The 10-sample 100 Gb marine dataset was created with CAMISIM from BIOM profiles of a deep-sea environment, using 155 newly sequenced marine isolate genomes from this environment and 622 genomes with matching taxonomic provenance from MarRef^79^, a manually curated database with completely sequenced marine genomes. Of these genomes, 303 (39.0%, 204 of database genomes (31.9%) and 99 new genomes (72.3%)) had a closely related strain present, with an ANI of 95% or more. Additionally, 200 newly sequenced circular elements including plasmids and viruses were added. For each sample, 5 Gb of paired-end short Illumina and long Nanopore reads were created (Supplementary Text).

The 100-sample 400 Gb strain madness dataset includes 408 newly sequenced genomes, of which 97% (395) had a closely related strain. For each sample, 2 Gb of paired-end short and long-read sequences were generated with CAMISIM, respectively, using the same parameters and error profiles as in CAMI 1^6^ (Supplementary Text).

The 21-sample 315 Gb plant-associated dataset includes 894 genomes. Of these, 224 are from the proGenomes^80^ terrestrial representative genomes, 216 are newly sequenced genomes from an *Arabidopsis thaliana* root rhizosphere, 55 are fungal genomes associated with the rhizosphere^81^, 398 are plasmids or circular elements and one *Arabidopsis thaliana* genome. 15.3% (137) of these genomes have at least one closely related genome present. For each sample, 5 Gb of paired-end short-read sequences, as well as 2x5 Gb long-read sequences mimicking Pacific Biosciences and Oxford Nanopore sequencing data, respectively, were generated. 90% of metagenome sequence data originate from bacterial genomes, 9% are fungal genome sequences, and 1% is from *A. thaliana*. To evaluate the assembly quality of single-sample versus cross-assembly strategies, 23 new genomes from eight clusters of closely related genomes were selected and added to the dataset in certain samples with predetermined abundances. For all three datasets, we generated gold standards for every metagenome sample individually and for the pooled samples, which included assemblies for short, long, and hybrid reads, genome bin and taxon bin assignments, and taxonomic profiles.

Finally, a 688 Mb paired-end Miseq metagenomic sequencing dataset of a blood sample from a patient with hemorrhagic fever was provided. Previous analysis of the sample had revealed sequences matching the genome of Crimean–Congo hemorrhagic fever orthonairovirus (CCHFV), and the presence of the viral genome was subsequently confirmed via PCR (with a Ct value of 27.4). To create a realistic dataset and case for the challenge while protecting the identity of the patient, a clinical case description derived from the true anamnesis and modified in ways consistent with the causative agent was created. Additionally, all reads mapping to the human genome were replaced by sequences from the same genomic regions randomly drawn from the 1000 genomes dataset^82^. Challenge participants were asked to identify the causal pathogen as well as all other pathogens present in the sample.

### Challenge organization

The second round of CAMI challenges assessed software for metagenome assembly, genome binning, taxonomic binning, and taxonomic profiling. In addition, a diagnostic pathogen prediction challenge was provided. As before, two metagenome “practice” benchmark datasets were created from public genomes and provided together with the standard of truth before the challenges, to enable contest participants to familiarize themselves with data types and formats. These included a 49-sample dataset modelled from Human Microbiome data^43^ and a 64-sample dataset modelled in taxonomic composition from mouse gut samples^83, 84^, with 5 Gb long (Pacific Biosciences, variable length with a mean of 3000 bp) and 5 Gb short (Illumina HiSeq2000, 150 bp) paired-end read sequences, respectively. Read profiles (read length and error rates) were created from sequencing runs on the MBARC-26 dataset^85^. Reference data collections with NCBI RefSeq, nr/nt and taxonomy from January 8th of 2019 were provided to participants, for use with reference-based methods in the challenges. For future benchmarking, use of these resources will facilitate method performance comparisons, as all genomes incorporated into the CAMI challenge datasets will be submitted to public sequence repositories.

The second CAMI challenge started on January 16th of 2019 (https://www.microbiome-cosi.org/cami/cami/cami2). Participants registered for download of the challenge datasets, with 332 teams registering from that time until January 2021. For reproducibility, participants could submit either a Docker container containing the complete workflow, a bioconda script or a software repository with detailed installation instructions, specifying all parameter settings and reference databases used. Assembly results could be submitted for short-read data, long-read data, or both data types combined. For methods incapable of submitting a cross-sample assembly for the entire dataset, a cross-sample assembly for the first ten samples of a dataset could be submitted. Participants could also submit single-sample assemblies for each of the first five samples of a dataset. Specification of the performance criteria for strain-aware assembly can be found in the Supplementary Material. The assembly challenge closed on May 17, 2019. Immediately afterwards, gold standard and MEGAHIT^31^ assemblies were provided for both datasets. The gold standard assemblies include all sequences of the reference genomes and circular elements covered by one short read in the combined metagenome datasets. Analysis of gold standard assembly binnings allowed us to assess binning performances independently of assembly quality. We assessed the contributions of assembly quality by comparing with the binning results on MEGAHIT assemblies. Profiling results were submitted for all individual samples and for the entire datasets, respectively. Binning results included genome or taxon bin assignments for analyzed reads or contigs of the provided assemblies for every sample of a dataset. Results for the pathogen detection challenge included predictions of all pathogens and a causal pathogen responsible for the symptoms outlined in a clinical case description provided together with the clinical metagenome dataset. The CAMI II challenges ended on October 25, 2019. Subsequently, another round of challenges (“CAMI II b”) on plant-associated data was offered starting on February 14, 2020. This closed on September 29, 2020 for assembly submissions and on January 31, 2021 for genome and taxonomic binning, as well as profiling.

Altogether 5,002 submissions of 76 programs were received for the four challenge datasets, from 30 external teams and CAMI developers (Supplementary Table 2). All genome data used for generation of the benchmark datasets as well as their metadata was kept confidential during the challenge and released afterwards (10.4126/FRL01- 006421672). To support an unbiased assessment, program submissions were represented with anonymous names in the portal (known only to submitters), and a second set of anonymous names for evaluation and discussion in the evaluation workshop, such that identities were unknown to all except for data analysis team (F.M., Z-L.D., A.F., A.S.), and program identities revealed only after a first consensus was reached.

### Evaluation metrics

In the following, we briefly outline the metrics used to evaluate the four software categories. For details, the reader is also referred to^20, 53, 61^.

#### Assemblies

Assemblies were evaluated with metaQUAST 5.1.0rc using the *--unique-mapping* flag. This flag allows every contig to be mapped at only a single reference genome position. In evaluation, we focused on commonly used assembly metrics such genome fraction, mismatches per 100 kb, duplication ratio, NGA50 and the number of misassemblies. The genome fraction specifies the percentage of reference bases covered by assembled contigs obtained by similarity-based mapping. Mismatches per 100 kb specify the number of mismatched bases in the contig-reference alignment. The duplication ratio is defined as the total number of aligned bases divided by genome fraction multiplied with reference length. NGA50 is a metric for measuring the contiguity of an assembly. For each reference genome, all contigs aligned to it are sorted by size and the NGA50 for that genome is defined as the length of the contig cumulatively surpassing 50% genome fraction. If a genome is not covered to 50%, NGA50 is undefined. Since we report the average NGA50 over all genomes, it was set to 0 for genomes with less than 50% genome fraction. Finally, the number of misassemblies describes the number of contigs which either contain a gap of more than 1kb, contain inserts of more than 1kb or align to different genomes. In addition to these metrics, we determined the strain recall and strain precision, similar to^38^, to quantify the presence of high-quality, strain-resolved assemblies. Strain recall is defined as the fraction of high-quality (more than 90% genome fraction and less than 100 mismatches per 100 kb) genome assemblies recovered for all ground truth genomes. Strain precision specifies the fraction of high-quality assemblies among all high genome fraction (more than 90%) assemblies.

#### Genome binning

For every predicted genome bin *b*, the true positives *TP*_*b*_ are the number of base pairs of the most abundant genome *g* in *b*, the false positives *FP*_*b*_ are the number of base pairs in *b* belonging to genomes other than *g*, and the false negatives *FN*_b_ are the number of base pairs belonging to *g* that are not in *b*.

Purity is defined for each predicted genome bin *b* as:

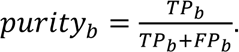

The average purity is a simple average of the purity of bins *b* in the set of all predicted genome bins *B*, that is:

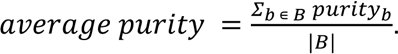

Completeness is defined for each genome *g* based on its mapping to a genome bin *b* that it is most abundant in, as:

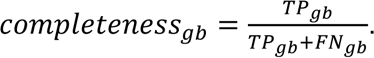

The average completeness is defined over all genomes in the sample, including those that are the most abundant in none of the predicted genome bins. Let *X* be the set of such genomes. The average completeness is then defined as:

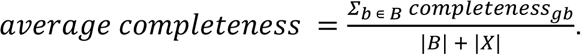

As another metric, we consider the number of predicted genome bins that fulfill specific criteria. Bins with >50% completeness and <10% contamination are denoted as “moderate or higher” quality bins and bins with completeness >90% and contamination <5% as high-quality genome bins, similarly as in CheckM^74^.

The Adjusted Rand index (ARI) is defined as in ^53^. The Rand index compares two clusterings of the same set of items. Assuming the items are base pairs of different sequences, base pairs belonging to the same genome that were binned together in the same genome bin are considered true positives, and base pairs belonging to different genomes that were put into different genome bins are considered true negatives. The Rand index is the sum of true positives and negatives, divided by the total number of base pairs. The ARI is a normalized variant of the Rand index, such that the result ranges between 1 (best) and 0 (worst; see ^53^ for a complete definition). As binning methods may leave a portion of the data unbinned, but the ARI is not suitable for datasets that are only partially assigned, it is computed for the binned portion only and interpreted together with the percentage of binned base pairs of a dataset.

#### Taxonomic binning

Metrics are calculated for each of the major taxonomic ranks, from superkingdom or domain to species. Purity and completeness for each taxonomic bin *b* (i.e., group of sequences and base pairs therein assigned to the same taxon) are computed by setting *TP*_*b*_to the number of base pairs of the true taxon *t* assigned to *b*, *FP*_*b*_ the number of base pairs assigned to *b* belonging to other taxa, and *FN*_b_ the number of base pairs of *t* not assigned to *b*. The average purity at a certain taxonomic rank is a simple average of the purity of all predicted taxon bins at that taxonomic rank.

The average completeness at a certain taxonomic rank is the sum of the completeness over all predicted taxon bins divided by the number of taxa *GS* in the gold standard at that taxonomic rank. That is:

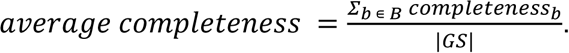

The accuracy at a certain taxonomic rank is defined as:

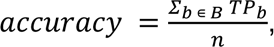

where *B* is the set of predicted taxon bins at that taxonomic rank and *n* is the total number of base pairs in *GS* for that taxonomic rank.

Average purity, completeness, and accuracy are also computed for a filtered subset *B*_*f*_ of *B* of each taxonomic rank, without the 1% smallest bins, and are denoted below *average purity*_*f*_, *average completeness*_*f*_, *accuracy*_*f*_ *B*_*f*_ is obtained by sorting all bins in *B* by increasing size in base pairs and filtering out the first bins whose cumulative size sum is smaller or equal to 1% of summed size of all bins in *B*. These metrics are then computed as:

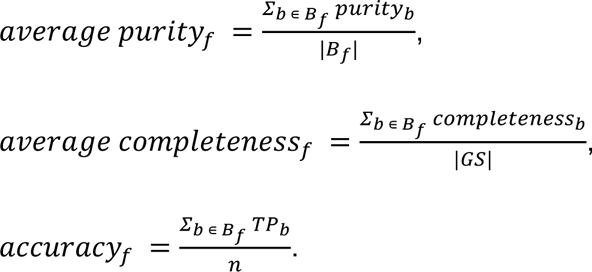

#### Taxonomic profiling

We determined purity and completeness in taxon identification, L1 norm and weighted UniFrac ^86^ as abundance metrics, and alpha diversity estimates using the Shannon equitability index, as outlined below. We also calculated the following summary statistic: for each metric, we ranked the profilers by their average performance over samples. Each was assigned a score for its ranking (0 for first place among all tools at a particular taxonomic rank, 1 for second place, etc.). These scores were then added over the taxonomic ranks, from domain to species, to give an overall summary ranking score.

The purity and completeness for a taxonomic profile measure a method’s ability to determine the presence and absence of taxa in a sample, at a certain taxonomic rank, without considering their relative abundances. Let the true positives *TP* and false positives *FP* be the number of correctly and incorrectly detected taxa, that is, taxa present or absent in the gold standard profile, respectively, for a certain sample and rank. Further, let the false negatives *FN* be the number of taxa that are in the gold standard profile but a method failed to detect. Purity, completeness, and F1-score are then defined as above.

The L1 norm error, Bray-Curtis distance, and weighted UniFrac error measure a method’s ability to determine the relative abundances of taxa in a sample. Except for the UniFrac metric (which is rank independent), these are defined at each taxonomic rank. Let *x*_*t*_ and *B*^∗^ be the true and predicted relative abundances of taxon *t* in a sample, respectively. The L1 norm gives the total error between *x*_*t*_ and *x\**_*t*_ in a sample, for all true and predicted *t* at a certain rank and ranges between 0 and 2. It is determined as:

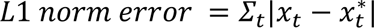

The Bray-Curtis distance is the L1 norm error divided by the sum of all abundances *x*_*t*_ and *x\**_*t*_ at the respective rank, that is:

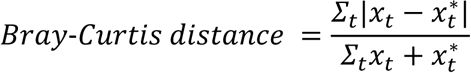

The Bray-Curtis distance ranges between 0 and 1. As the gold standards usually contain abundances for 100% of the data, it is equal to half of the L1 norm error if the profiler made predictions also for 100% of the data, and higher otherwise.

The weighted UniFrac metric uses differences between predicted and actual abundances weighted by distance in the taxonomic tree. It ranges between 0 (best) and 16 (worst). We use the EMDUnifrac implementation of the UniFrac distance ^87^.

The Shannon equitability index is defined for each rank as:

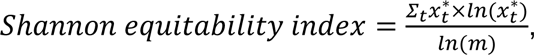

where *m* is the total number of taxa *t*. The index ranges from 0 to 1, with 1 indicating complete evenness. As the diversity estimate is computed from a predicted profile alone, we assess its absolute difference to the index of the gold standard for comparison.

### Data availability and accession code availability statements

A Life Science Reporting Summary for this paper will be made available. The benchmarking challenge and exemplary datasets (for developers to familiarize upfront with data types and formats) are available in PUBLISSO with the DOIs 10.4126/FRL01- 006425521 (marine, strain madness, plant-associated), 10.4126/FRL01-006421672 (mouse gut), and 10.4126/FRL01-006425518 (human) and on the CAMI data portal for download (https://data.cami-challenge.org/participate). Datasets include gold standards, assembled genomes underlying benchmark data creation, NCBI taxonomy versions, and reference sequence collections for NCBI RefSeq, nt and nr (status 019/01/08). Benchmarked software outputs are available on Zenodo (https://zenodo.org/communities/cami/). Further software and scripts used for data analyses, and results are available at https://github.com/CAMI-challenge/second_challenge_evaluation. Supplementary Table 2 specifies the evaluated programs, parameters used, and installations options, including software repositories, Bioconda package recipes, Docker images, Bioboxes, and Biocontainers. Source data and scripts for Figures 1-5 are available online (https://github.com/CAMI-challenge/second_challenge_evaluation/).

## Acknowledgements

The authors thank all members of the metagenomics community who provided inputs and feedback on the project in public workshops and gratefully acknowledge funding of the DZIF (Project number TI 12.002_00; F.Mey.), German Excellence Cluster RESIST (EXC 2155 – Projektnummer 390874280; Z-.L.D.). D.K. was supported in part by the National Science Foundation under Grant No. 1664803; A.G. by St. Petersburg State University (grant ID PURE 73023672); D.A., A.Kor., D.M., and S.N. by the Russian Science Foundation (grant 19-14-00172); C.T.B. and L.I. in part by the Gordon and Betty Moore Foundation’s Data-Driven Discovery Initiative through Grants GBMF4551 to C.T.B.; R.C. and R.V. by ANR Inception (ANR-16-CONV-0005) and PRAIRIE (ANR-19-P3IA-0001); S.D.K. by the European Research Council (ERC) under the European Union’s Horizon 2020 research and innovation programme (ERC-COG-2018); J.K. and E.R.R. by the National Science Foundation under Grant No. 1845890; S.M. partially by National Science Foundation grants 2041984; V.R.M. by the Tony Basten Fellowship, Sydney Medical School Foundation. G.L.R. and Z.Z. partially by NSF grants 1936791 and 1919691; M.T. by the European Research Council (ERC) under the European Union’s Horizon 2020 research and innovation programme (ERC-COG-2018); S.Z. by the Shanghai Municipal Science and Technology Commission [2018SHZDZX01], 111 Project [B18015]; S.Ha. by the Deutsche Forschungsgemeinschaft (DFG, German Research Foundation) through the ‘2125 DECRyPT’ Priority Programme; R.E., E.Go., Zho.W., and A.T. by the DOE Office of Biological and Environmental Research under contract number DE-AC02-05CH11231. This research used resources of the National Energy Research Scientific Computing Center, which is supported by the Office of Science of the U.S. Department of Energy under Contract No. DE-AC02-05CH11231. The work conducted by the U.S. Department of Energy Joint Genome Institute, a DOE Office of Science User Facility, is supported under Contract No. DE-AC02-05CH11231.

## Author contributions

F.Mey., A.F., Z.-L.D., D.K., M.A., D.A., F.B., D.B., J.J.B., C.T.B., J.B., A.Bu., B.C., R.C., P.T.L.C.C., A.C., R.E., E.E., E.Ge., E.Go., S.Ho., P.H., L.I., H.J., S.D.K., M.K., A.Kor., J.K., N.L., C.Le., C.Li., A.L., F.M.-M., S.M., V.R.M., C.M., P.M., D.M., D.R.M., A.M., N.N., J.N., S.N., L.O., L.P., P.P., V.C.P., J.S.P., S.Ras., E.R.R., K.R., B.R., G.L.R., H.-J.R., V.S., N.Se., E.S., L.S., F.S., S.S., A.T., C.T., M.T., J.T., G.U., Zho.W., Zi.W., Zhe.W., A.W., K.Y., R.Y., G.Z., Z.Z., S.Z., J.Z., and A.S. participated in challenge and created results; P.W.D., L.H.H., T.S.J., T.K., A.Kol., E.M.R., S.J.S., N.P.W., R.G.-O., P.G., S.Ha., S.Hä., A.Kh., F.Ma., F.Mes., S.Rad., P.S.-L., N.Sm., and T.S. generated and contributed data; A.F., A.Br., A.S., and A.C.M. generated benchmark datasets; F.Mey., D.K., A.G., M.A.G., L.I., G.L.R., Z.Z., and A.C.M. implemented benchmarking metrics; F.Mey., A.F., D.K., A.S., Z.-L.D., and A.C.M. performed evaluations and interpreted results with comments from many authors; F.Mey., A.F., Z.-L.D., D.K., A.G., G.R., F.B., R.C., P.W.D., A.E.D., R.E., D.R.M., A.M., E.R.R., B.R., G.L.R., H.-J.R., S.S., R.V., Z.Z., A.Br., A.S., and A.C.M made conceptual inputs to challenge design or evaluation; F.Mey., A.C.M., A.F., D.K. and Z.-L.D. wrote the paper with comments from many authors; A.S. and A.C.M. conceived the research with input from many authors.

