## Supplementary information for "Critical Assessment of Metagenome Interpretation - the second round of challenges"

### CAMI II benchmark data generation

#### Marine and plant-associated datasets

To generate the CAMI II marine dataset, we used the CAMISIM[1] simulator to simulate a hybrid long and short-read shotgun metagenome dataset, including ten samples with a taxonomic composition matching an arctic marine microbiome[2] profile. The taxonomic profile was generated using USEARCH[3] for OTU clustering and taxonomic classification with the RDP[4] classifier based on pyrosequencing 16S data from an arctic marine environment (ERP003605). As input genomic data, 622 high quality genomes from the MarRef database[5] as well as 224 representative terrestrial genomes from proGenomes[6] were used, as well as 176 and 216 novel genomes from the respective environment as well as 598 short circular elements currently not in the public domain. The new genomes were assembled with the SPAdes[7] assembler, version 3.12, using the --careful flag and default arguments otherwise. For each genome, sequences of length 1 kb or less were removed and a taxonomic annotation with CAMITAX[8] was performed. CAMITAX includes a CheckM[9] run, whose contamination and completion values were used to remove the new as well as multi-contig MarRef genomes, if completeness was less than 90% or contamination higher than 5%.

For the database genomes, where a taxonomic classification was available, CAMITAX was used as a consistency check, and the lowest common ancestor of the CAMITAX classification and the original taxonomic classification was chosen. The resulting taxonomically annotated genome set (including the newly sequenced genomes) and their taxonomic classification along with a 16S rRNA profile were then used to generate shotgun metagenome samples with CAMISIM in the community design mode. Within CAMISIM, a community genome abundance profile was generated by mapping the taxa from 16S rRNA profile to the input genomes and abundances were assigned accordingly. Genomes not mapped to any taxa were subsequently randomly assigned to the most

abundant, non-assigned OTU, such that all input genomes were included in the dataset. In the next step, plasmids were added to the generated community genome abundance profile. We used plasmids specifically sequenced and classified for CAMI. Since the plasmids are expected to be circular while fasta files only allow a linear representation, the original fasta files were treated in the following way. Given a plasmid sequence of *n* bases, split the sequence in 10 equally long segments, *n_1, …, n_10*, where each *n_i* consists of the *i-th tenth* bases of the input sequence. Then create 10 permutations of the input sequence, where always the first segment is moved to the end of the previous permutation. When the original sequence *n_1, …, n_10* is the first permutation, then *n_2, …, n_10, n_1* is the second, *n_3, …, n_10, n_1, n_2* the third and so forth.
The plasmids were identified as either virus or plasmid (or unknown) using the following process[10]:

- Prodigal[11] complete gene prediction (including genes overlapping sequence ends)
- Annotation using hmmscan[12] (max e-value 1e-4) and PFAMv27[13]
- Any sequence with plasmid replication, mobilization, or stability was classified as plasmid
- Any sequence without these plasmid markers, but with a viral replication or capsid gene was classified as virus/phage.

Any sequence with neither of the features was designated unknown.
To add the plasmids treated this way, the desired number, 200 for the marine challenge, were selected and randomly assigned to the 200 highest abundant genomes to emulate the affiliation of the plasmids with the genomes. Since plasmids are highly abundant in metagenomic datasets, they were chosen to have ~15x the abundance of the input genomes, in particular this meant that every permutation was assigned roughly 1.5x of the affiliated genomes' abundance. The exact number was calculated by the formula: ab_plasmid = ab_genome * 1.5 * N(1,0.1).
Given all the genomes and plasmids with their respective abundance, CAMISIM could finally be run to create the challenge dataset.
The scripts and command line options are provided on Github under https://github.com/CAMI-challenge/second_challenge_evaluation

**Strain madness dataset**

For the strain madness dataset, 408 newly genomes were sequenced, assembled, and taxonomically classified using CAMITAX. 395 have a closely related genome present with 180 *Streptococcus pneumoniae*, 97 *Escherichica coli*, 47 *Klebsiella pneumoniae*, 21 *Staphylococcus aureus*, 21 *Enterococcus faecium* and 27 further *Enterobacteriacaeae*. Additionally, 13 unique genomes from a mouse gut, mainly consisting of *Lactobacillus* and Bacteroides, were added. This dataset was simulated without a BIOM profile as input, but instead using the *differential* mode of CAMISIM.

### CAMI metagenome assembly evaluation for strain-specific assemblies

To assess strain-specific assemblies, we engaged the computational metagenomics community in the CAMI II evaluation workshop and defined assembly properties and evaluation strategies. Assessments in CAMI are done according to these agreed strategies, which have been incorporated in MetaQUAST version 5.1.0rc and are available via the new `--unique-mapping` option (http://cab.cc.spbu.ru/quast/manual.html#unique_mapping). The following command shows how to apply the new option on the strain madness coassembly, as an example.

Synopsis: `quast-5.1.0rc1/metaquast.py --reuse-combined-alignments --no-icarus -o ../strmgCAMI2_co_assembly_metaquast-5.1.0rc1 -r `cat ../refs` -t 28 --unique-mapping ../strmgCAMI2_pooled/GS_*`

We consider two genomes to be different strains of the same species if they have >95% average nucleotide identity.

A *consensus* (or *strain-unresolved*) assembly is one where each contig may correspond to 1 or many strains (*strain-unresolved* contig), and reciprocally (and importantly), any genomic region that has one or more homologs across different strains is represented in only one contig. Assemblers, e.g., MEGAHIT and metaSPAdes, that create such assemblies are *strain-oblivious* assemblers.

A *strain-resolved, or strain-specific* assembly is one where each contig either i) maps equally likely to >1 strains (*core* contigs), or ii) maps unambiguously to only 1 strain (*strain-specific* contigs). Assemblers that create such assemblies are strain-aware or strain-resolved assemblers. In addition, core contigs should be present in as many copies as there are genomes containing such regions (see example below).

#### Evaluation of strain-specific assemblies

Consider the following reference genomes:

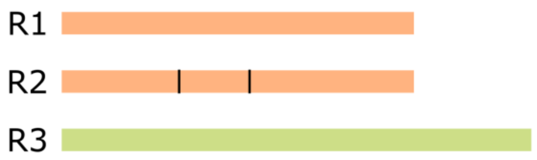


R1 and R2 are two strains of the same species (%-identity higher or identical to our threshold set above). R3 is a different species, without homologous regions with R1 and R2.

**Case 1: two (extreme) examples of assemblies**

Assembly **A1**

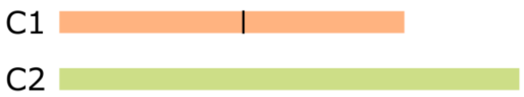


A1 is a consensus assembly. Here, contig C1 corresponds to a consensus of R1 and R2, and contig C2 corresponds exactly to R3.

Assembly **A2**

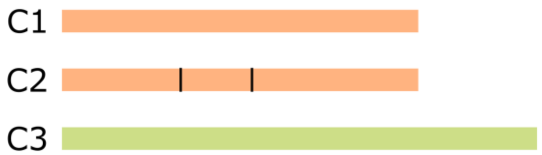


The contigs of A2 correspond exactly to the reference genomes. A2 is a strain-specific assembly. Contigs C1 and C2 are >= 95% identical.

Between **A1** and **A2**, in the context of strain-aware evaluation, we favor assembly **A2** over **A1**. In particular, recovered genome fraction will be higher for **A2** than for **A1**.

**Case 2: two more realistic assemblies**

**A3**

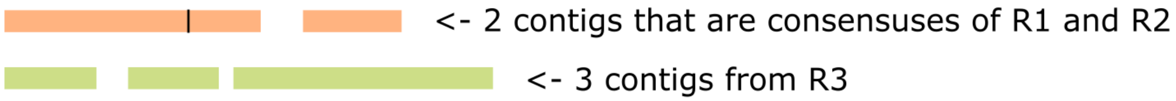


**A4**

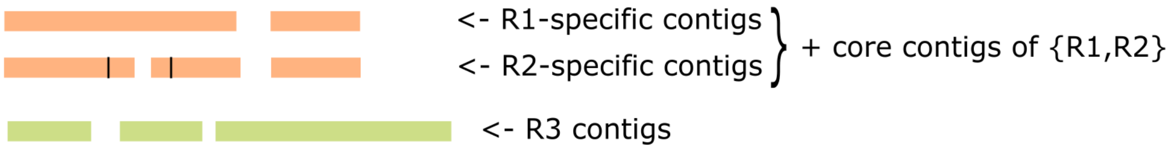


CAMI performed a strain-aware evaluation. Thus, in that context, we prefer assemblies like **A4** to assemblies like **A3** (despite the fragmentation in R2 contigs). We also prefer **A2** to **A4**.

### Supplementary figures

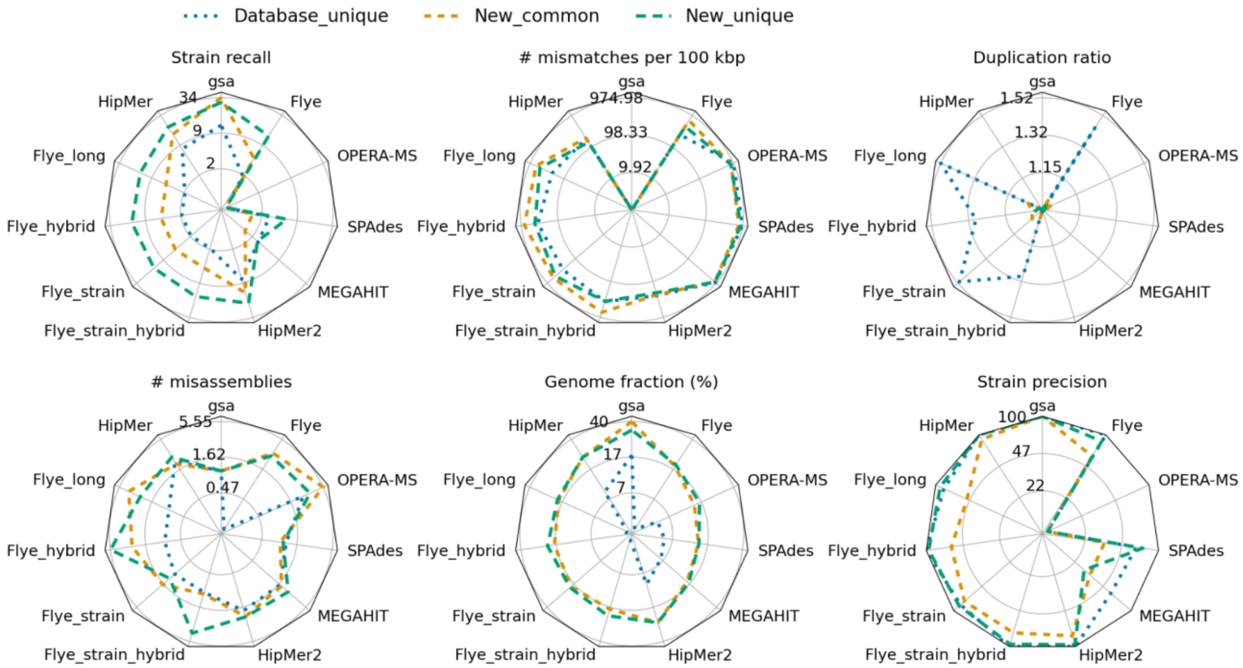


**Supplementary Fig. 1:** Radar plots of strain recall, mismatches per 100 kb, duplication ratio, misassemblies, genome fraction, and strain precision for assemblers on the plant-associated dataset.

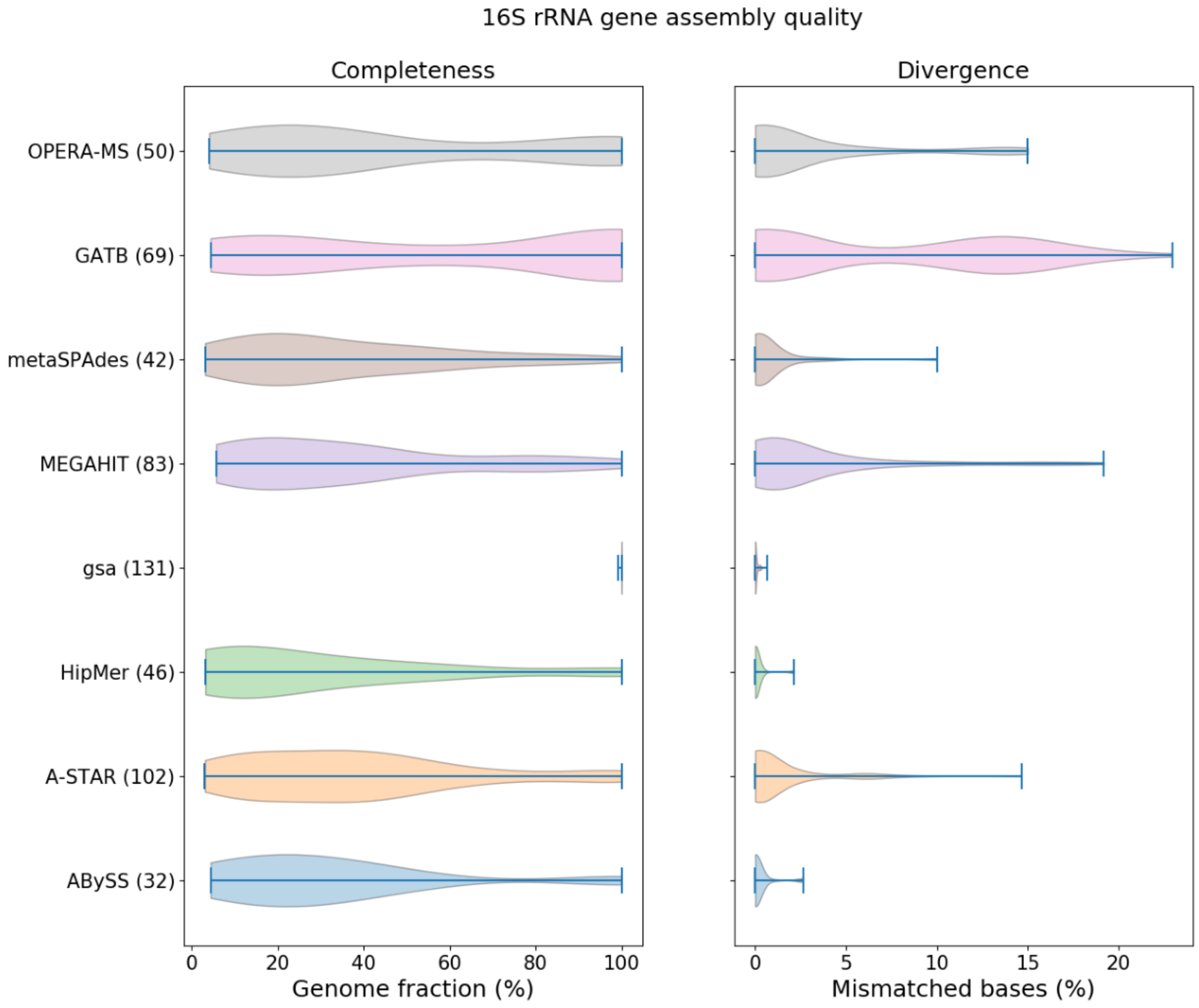


**Supplementary Fig. 2**: Completeness and divergence of assemblies on the 16S rRNA gene of 58 high-quality database genomes. Evaluated using metaquast against the 16S rRNA gene sequence of high-quality genomes, extracted from NCBI. To avoid mappings from other genomes, the contigs were aligned to the high-quality genomes first and each "contig bin" evaluated individually against the corresponding genome using QUAST[14]. Completeness describes the genome fraction, Divergence the number of mismatched bases, blue bars show the standard deviation. The number in brackets denoted the total number of reconstructed 16S rRNA sequences, i.e., the gold standard (denoted by gsa) contained 131 16S rRNA gene sequences in the 58 genomes and A-STAR recovered 102 of them.

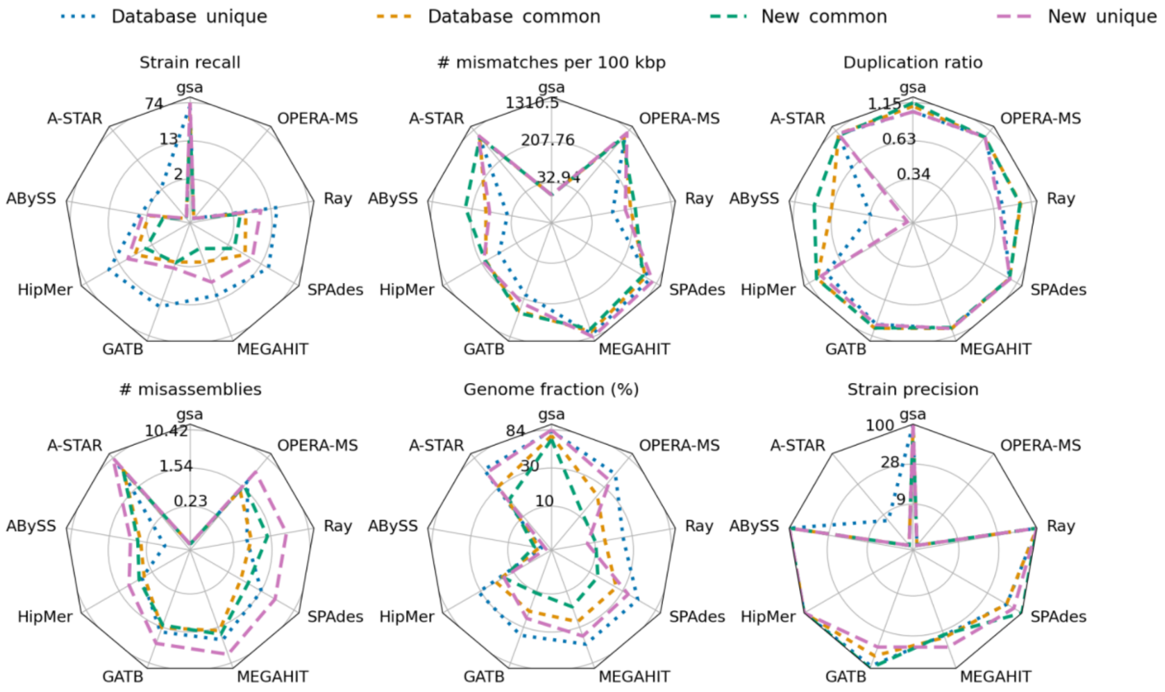


**Supplementary Fig. 3:** Radar plots comparing the assembly quality of new and database genomes and common and unique genomes of the marine dataset.

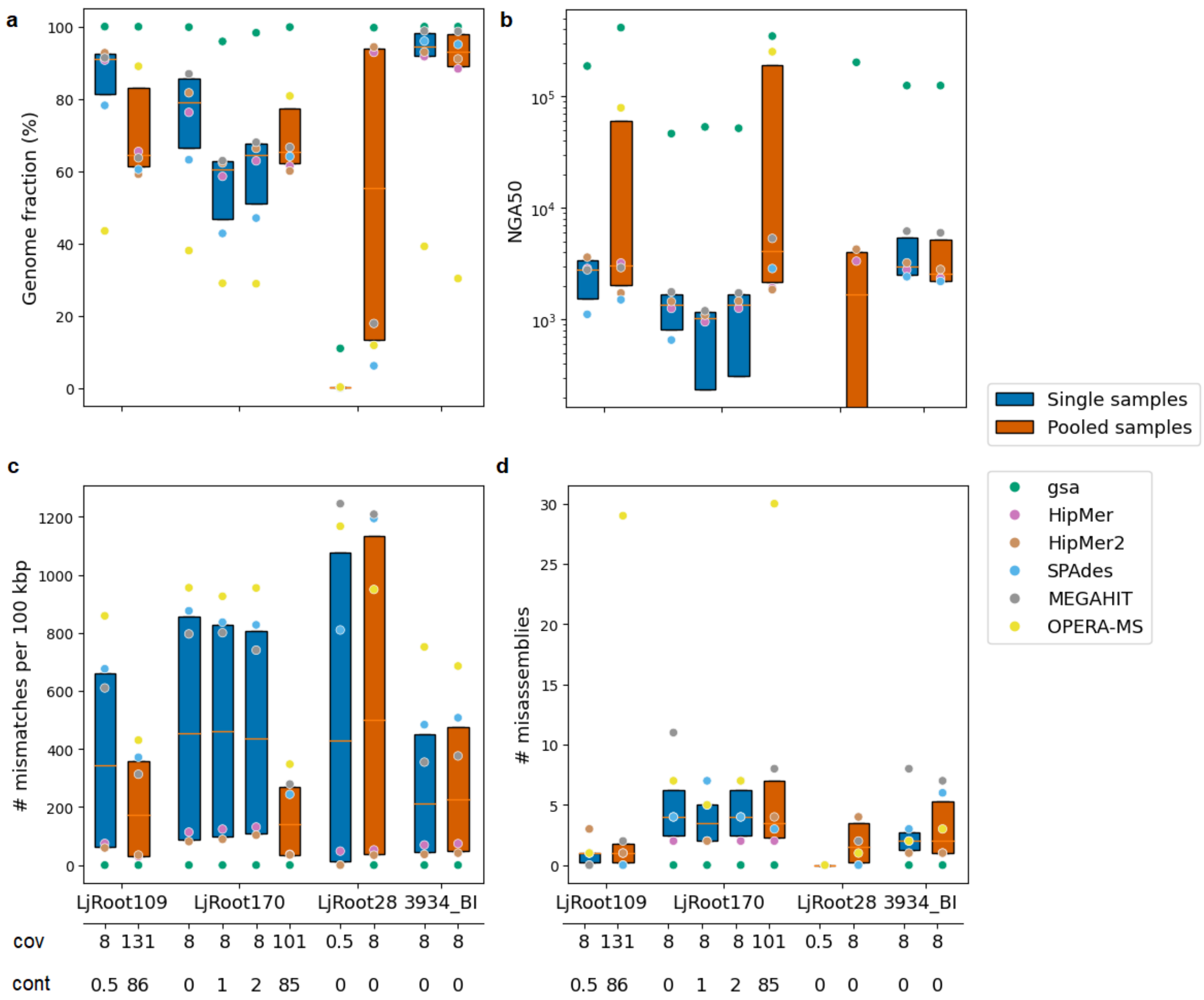


**Supplementary Fig. 4:** Boxplots of **(a)** genome fraction, **(b)** NGA50, **(c)** number of mismatches per 100 kb and **(d)** number of misassemblies of four spiked genomes for single (blue) and pooled samples (orange) of the plant-associated dataset. NGA50 is shown with log-scale; the individual assemblers for which single sample and pooled assemblies were available are color-coded; the gold standard (green) is denoted with gsa. “cov” denotes the coverage of the corresponding genome in the single or pooled samples and “cont” (contamination) the total coverage of closely related genomes present in the sample.

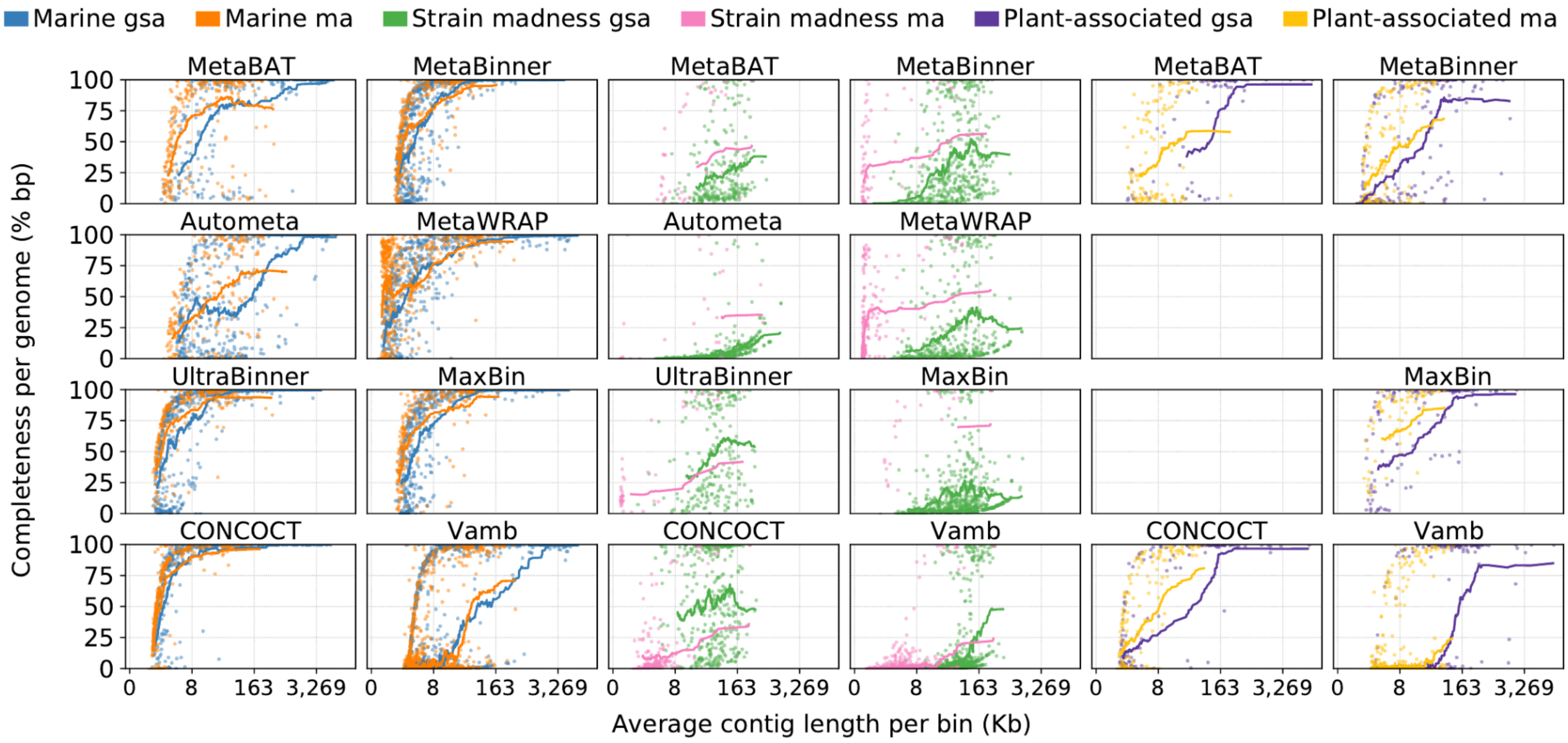


**Supplementary Fig. 5**: Average length of the contigs per genome bin vs. the completeness of the corresponding genome on different datasets per genome binner. "gsa" and "ma" are the binnings of the gold standard and MEGAHIT assemblies of a dataset, respectively. Lines give the running average completeness of 50 consecutive genomes ordered by the average contig length in the bins. For each method, the best-performing submission in terms of F1-score is shown.

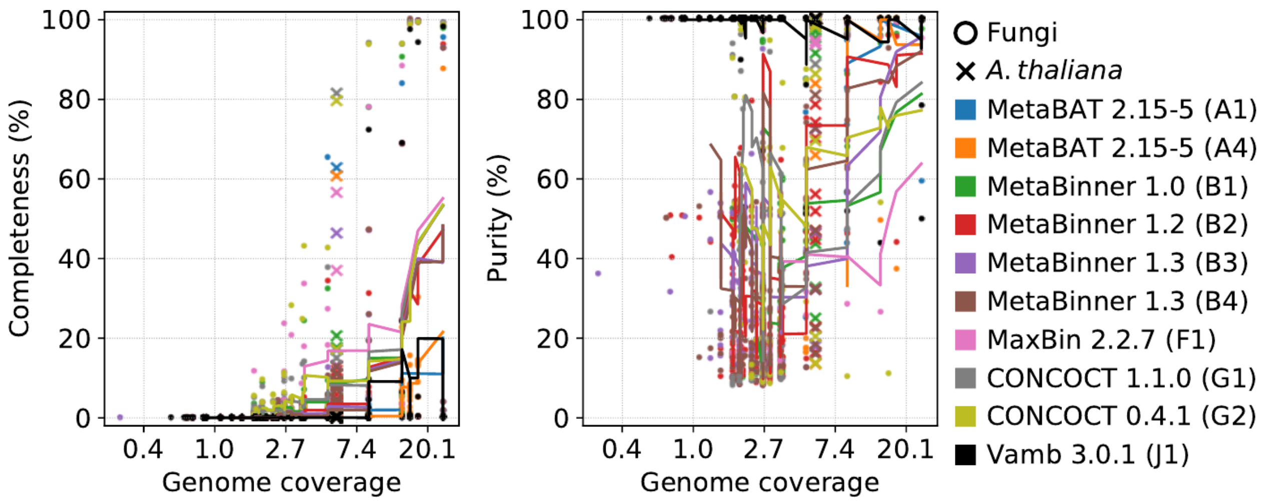


**Supplementary Fig. 6**: Coverage of fungal genomes (circles) and the *A. thaliana* genome (cross marks) in the plant-associated dataset recovered by genome binners vs. completeness per genome (left) and purity per bin (right). Lines indicate the running average completeness or purity of 10 consecutive fungal genomes or bins ordered by coverage. Raw data is available at https://github.com/CAMI-challenge/second_challenge_evaluation/blob/master/assembly/scripts/data/funghi_coverage.tsv.

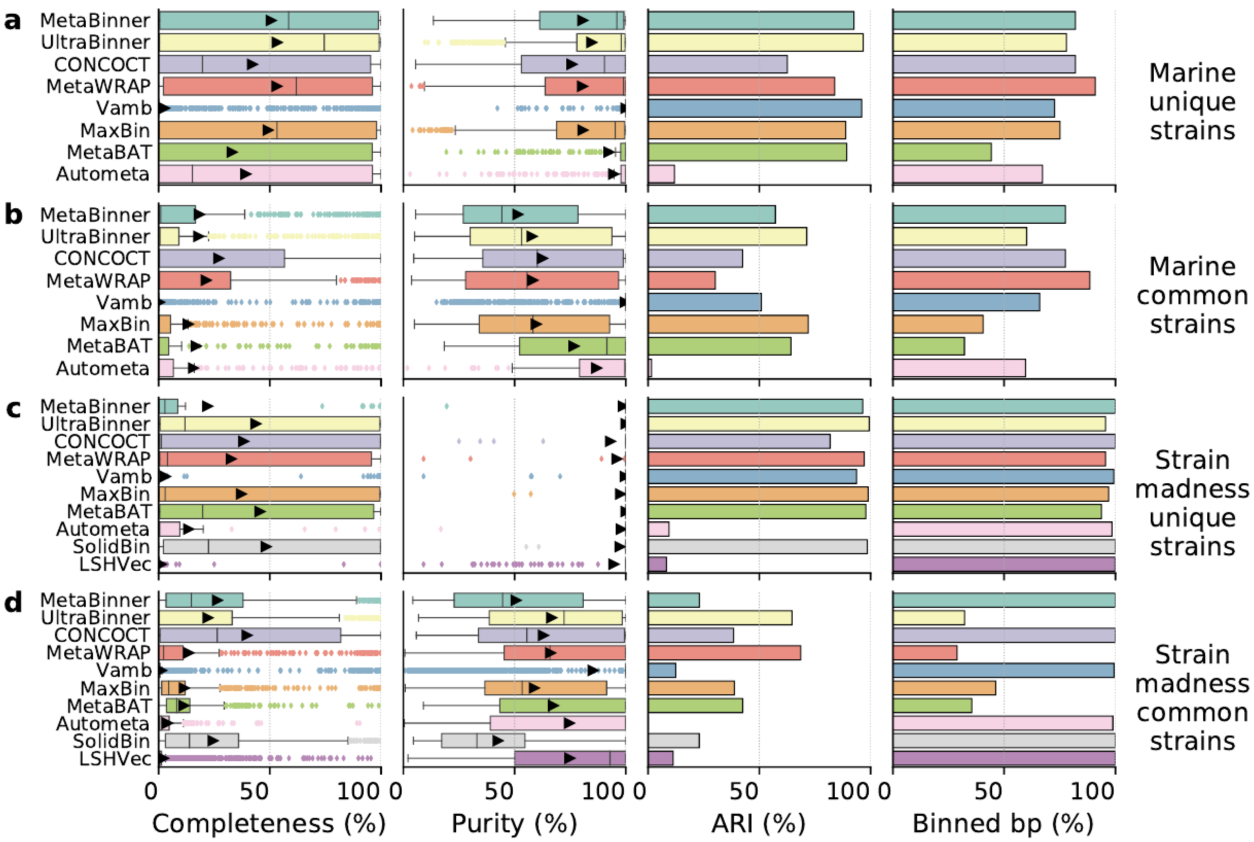


**Supplementary Fig. 7:** Effect of strain diversity on average completeness, purity, adjusted Rand index (ARI), and percentage of binned bp for genome binning of unique and common strains of the marine and strain madness short-read GSAs. Genomes were reconstructed by genome binners for genomes of unique strains with ANI < 95% to others and common strains with ANI ≥ 95% to each other.

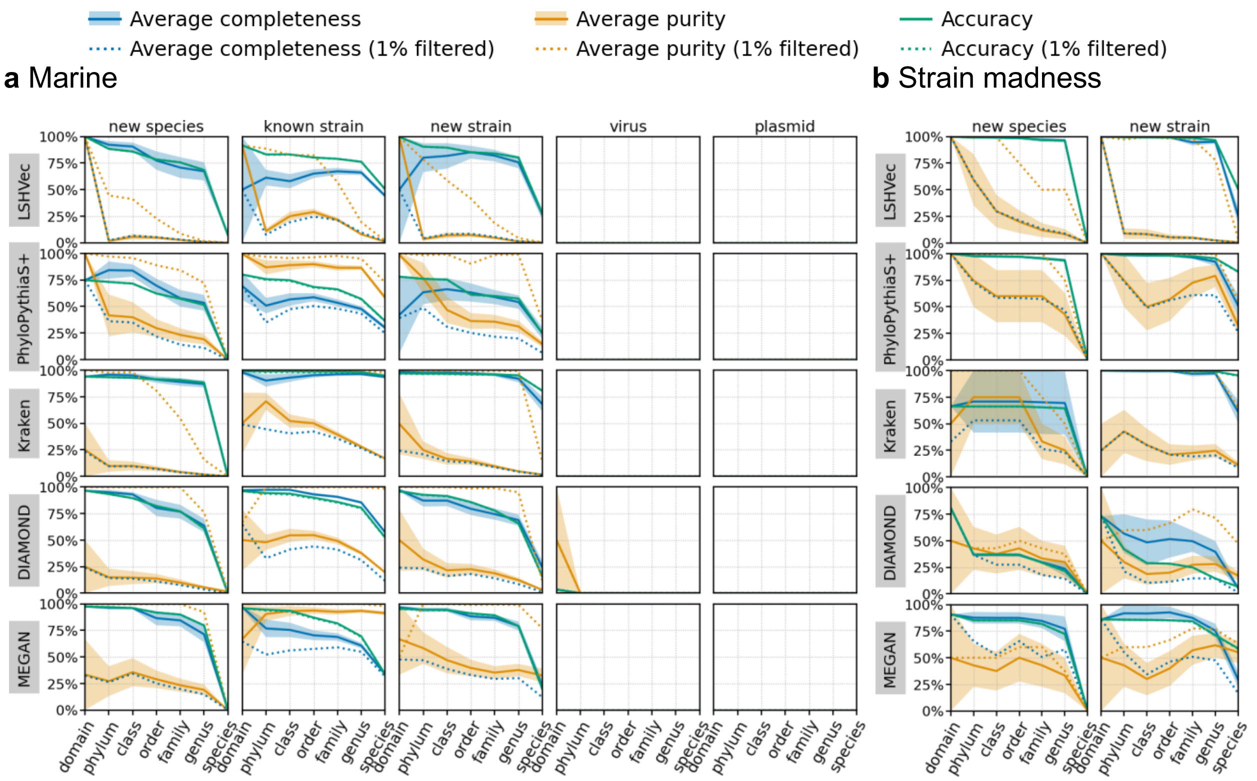


**Supplementary Fig. 8:** Taxonomic binning performance across ranks for the marine (a) and strain madness (b) datasets split by their taxonomic distances to public genomes as new species or strains, known species or known strains, viruses, or plasmids. A genome is classified as "new species" if no genome of that species is present in the NCBI RefSeq database. Metrics are computed over unfiltered and 1% filtered predicted bins (see main text). Shaded bands show the standard error across bins.

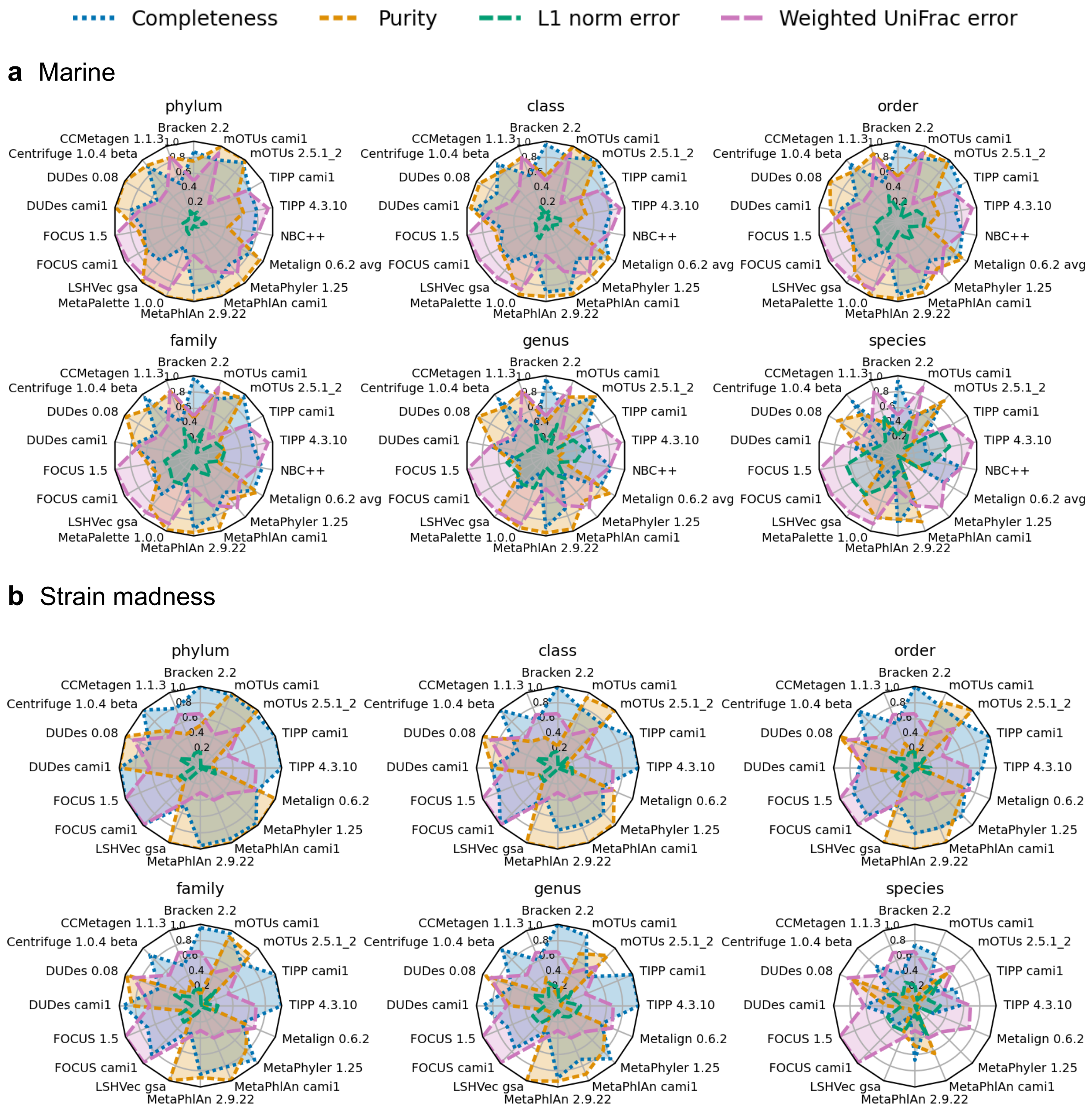


**Supplementary Fig. 9: Taxonomic profiling results for the marine (a) and strain madness (b) datasets from phylum to species ranks.**

L1 norm is divided by 2 to be in the range between 0 and 1, as completeness and purity. Weighted UniFrac error is normalized by the maximum value obtained by a method.

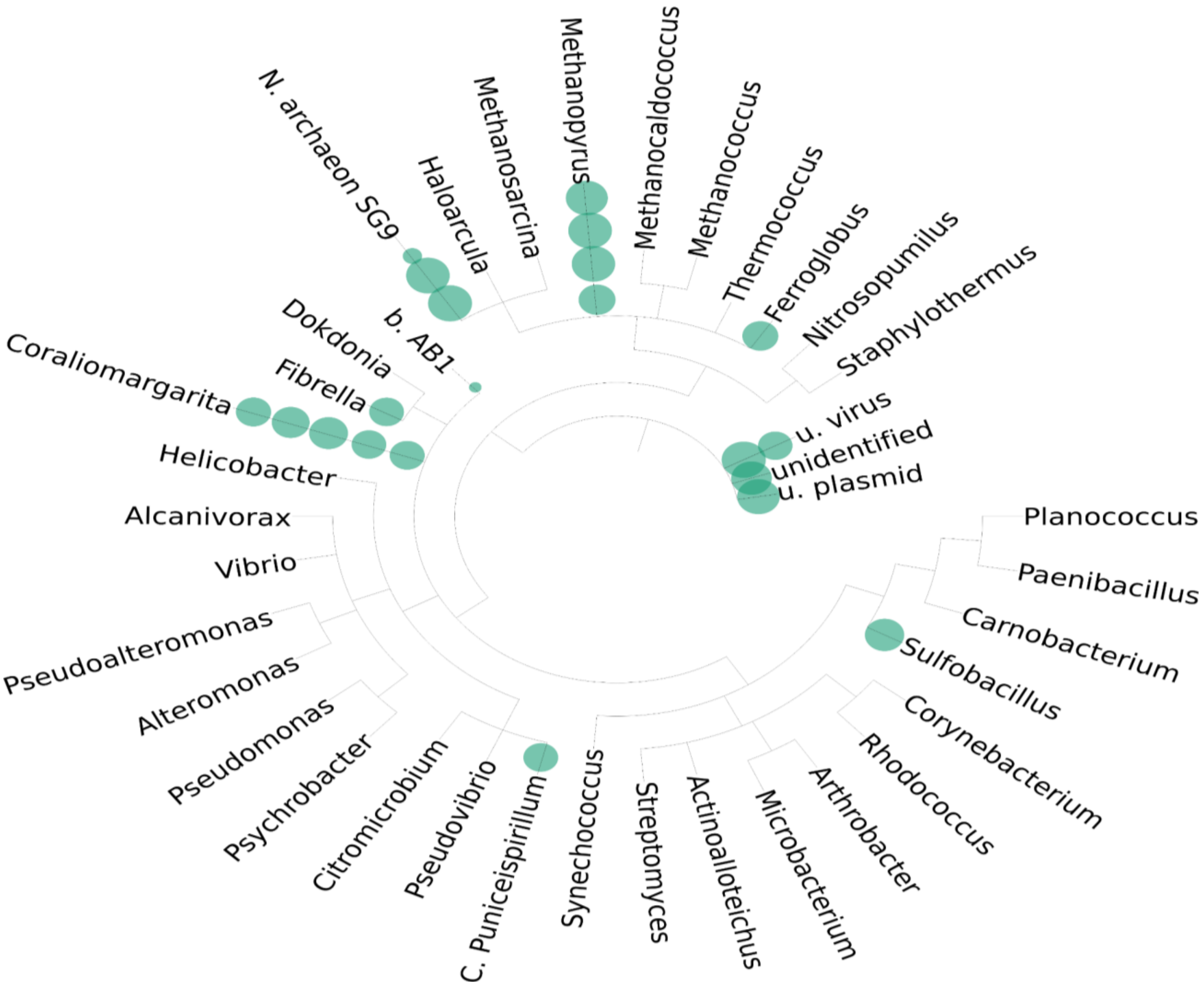


**Supplementary Fig. 10: Taxonomic tree depicting most difficult to detect taxa in the marine dataset.**

MeTEA (https://github.com/mag535/MeTEA) was utilized to identify the gold standard taxa most frequently missed by all tools. TAMPA (https://github.com/dkoslicki/TAMPA) was then used to depict these taxa on a tree. Green discs represent abundance in the ground truth, averaged over all marine datasets. Multiple discs represent different taxonomic ranks.

**a**

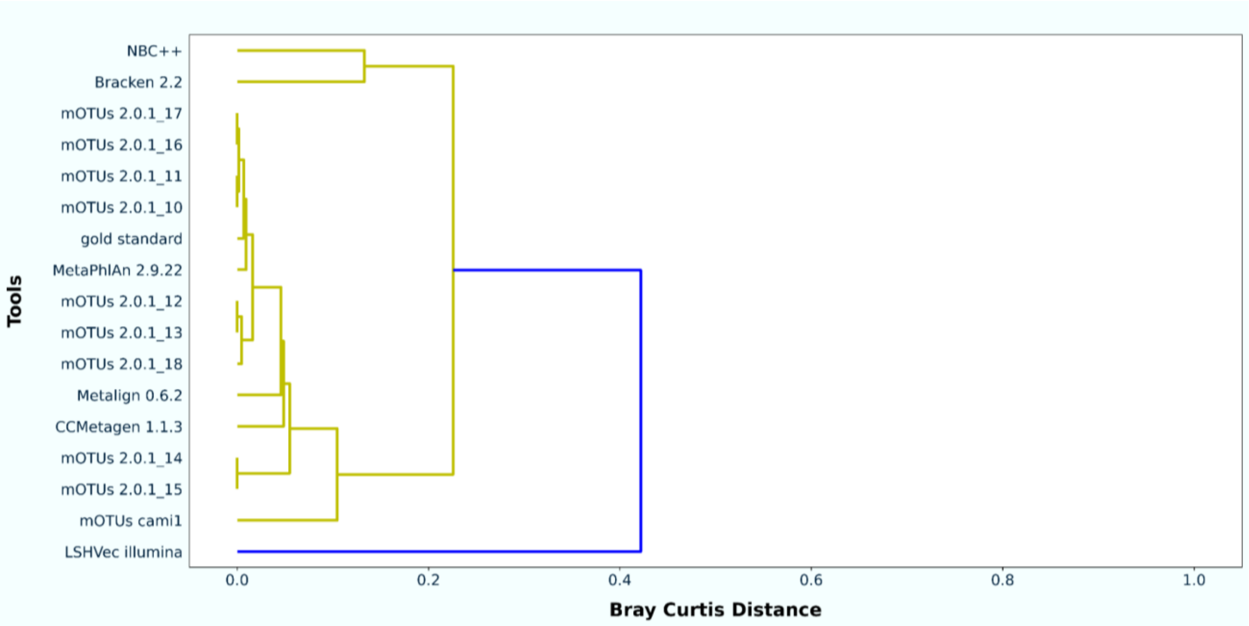


**b**

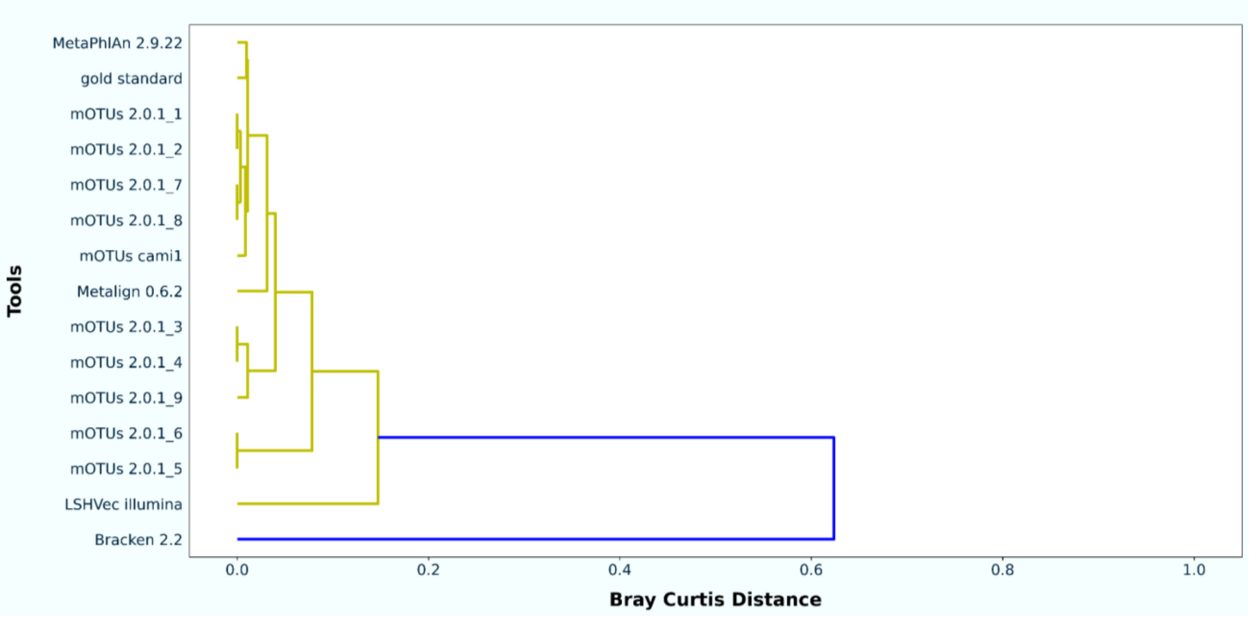


**Supplementary Fig. 11: Tool performance clustering.**

MeTEA (https://github.com/mag535/MeTEA) was utilized to cluster tool performance in the following way: at each taxonomic rank, MeTEA computes the F1 score for each tool and each taxa in the gold standard and then averaged over all ranks. The Bray-Curtis dissimilarity was used for the hierarchical clustering. Part **a** shows tool similarity for short-read marine dataset submissions and **b** depicts tool similarity for those that submitted results for the short-read strain madness dataset.

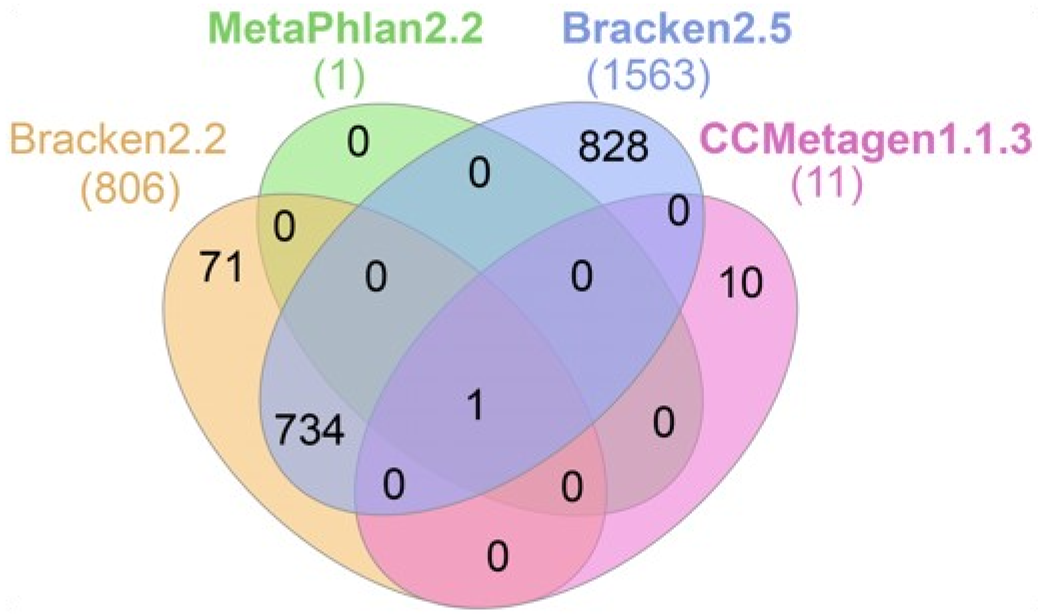


**Supplementary Fig. 12: Venn diagram of taxa predicted by different submissions to the clinical pathogen detection challenge.** Shown are methods that included the causal pathogen among the predicted taxa (total number in brackets) submitted to the challenge.
